# The Mechanosensitive Ion Channel MSL10 Modulates Susceptibility to *Pseudomonas syringae* in *Arabidopsis thaliana*

**DOI:** 10.1101/2021.08.18.456837

**Authors:** Debarati Basu, Jennette M. Codjoe, Kira M. Veley, Elizabeth S. Haswell

## Abstract

Plants sense and respond to molecular signals associated with the presence of pathogens and their virulence factors. Mechanical signals generated during pathogenic invasion may also be important, but their contributions have rarely been studied. Here we investigate the potential role of a mechanosensitive ion channel, MscS-Like (MSL)10, in defense against the bacterial pathogen *Pseudomonas syringae* in *Arabidopsis thaliana*. We previously showed that overexpression of MSL10-GFP, phospho-mimetic versions of MSL10, and the gain-of-function allele *msl10-3G* all produce dwarfing, spontaneous cell death, and the hyperaccumulation of reactive oxygen species. These phenotypes are shared by many autoimmune mutants and are frequently suppressed by growth at high temperature in those lines. Here, we found that the same was true for all three *MSL10* hypermorphs. In addition, we show that the SGT1/RAR1/HSP90 co-chaperone complex was required for dwarfing and ectopic cell death, PAD4 and SID2 were partially required, and the immune regulators EDS1 and NDR1 were dispensable. All *MSL10* hypermorphs exhibited reduced susceptibility to infection by *P. syringae* strain *Pto* DC3000, *Pto* DC3000 expressing the avirulence genes *avrRpt2* or *avrRpm1*, but not *Pto DC3000 hrpL*, and showed an accelerated induction of *PR1* expression compared to wild-type plants. Null *msl10-1* mutants were delayed in *PR1* induction and displayed modest susceptibility to infection by COR-deficient *Pst*. Finally, stomatal closure was reduced in *msl10-1* loss-of-function mutants in response to *Pst COR^−^*. These data show that MSL10 modulates pathogen responses and begin to address the possibility that mechanical signals are exploited by the plant for pathogen perception.

## INTRODUCTION

Plant immune receptors achieve a two-tiered response to pathogens (reviewed in (Chiang and Coaker 2015; Saijo et al. 2018; Jones et al. 2016)). The first line of defense relies on plasma membrane-localized pattern recognition receptors (PRRs), which facilitate molecular recognition of microbe- or damage-associated molecular patterns (MAMPs or DAMPs, respectively) and subsequently trigger pattern-triggered immunity, or PTI (reviewed in (Saijo et al. 2018)). However, pathogens evade PTI responses by delivering molecular virulence factors or effectors that subvert basal defenses into host cells (reviewed in (Toruño et al. 2015)).

To counter such pathogens, plants have in turn evolved cytosolic receptors which constitute the second layer of plant defense (reviewed in (Jones et al. 2016)). These intracellular receptors contain nucleotide-binding leucine-rich repeats and are thus known as NB-LRRs or NLRs. Both TIR-NLR and CC-NLRs detect the presence of intracellular effectors and activate effector-triggered immunity or ETI (reviewed in (Cui et al. 2015). Some NLRs recognize and bind effectors directly, while others recognize the host proteins that effectors modify (reviewed in (Khan et al. 2018)). It has recently been discovered that ZAR1 NLRs oligomerize to form a large complex that serves to detect the activity of several bacterial effectors and is capable of channeling calcium and triggering defense responses (Bi et al. 2021; Wang et al. 2019).

As described above, the molecular perception of pathogens and their effectors by PRRs and NLRs is central to the plant immune response. However, there is growing evidence that mechanical signals generated during invasion also elicit plant defense responses. For example, Arabidopsis, strawberry, and bean plants acquire transient immunity to necrotic or wounding pathogens following mild mechanical stimulus (reviewed in (Coutand 2020)). It is not yet known if pathogens create detectable mechanical perturbations, if the mechanoperception of pathogens utilizes the same pathways as defenses do, nor how such signals would be detected.

One mechanism by which the plant may recognize pathogen-associated mechanical signals is the activation of mechanosensitive (MS) ion channels. Multiple families of MS ion channels mediate the flux of ions across a membrane in response to increased membrane tension in plants (reviewed in (Basu and Haswell 2017; Guerringue et al. 2018)). Genetic and biochemical data support roles for several proteins that are homologs of known MS ion channels in the response to pathogens. Plants lacking the AtPIEZO channel are unable to fully suppress long-distance movement of viruses (Zhang et al. 2019), and OSCA1.3 is required for immunity-based stomatal closure (Thor et al. 2020). The putative MS ion channel MscS-Like (MSL) 4 contributes to basal immunity and MAMP-induced responses (Zhang et al. 2017).

MSL10 is a ubiquitously expressed homolog of MSL4 and is known to form a mechanically activated ion channel both in root cells and in Xenopus oocytes (Maksaev and Haswell 2012; Haswell et al. 2008) making it an intriguing candidate for a sensor of pathogen invasion. It is directly activated through membrane tension and is largely non-selective, with a slight preference for anions (Maksaev and Haswell 2012). Based on homology to other MscS-like channels, the MSL10 channel is likely a homoheptamer (Bass et al. 2002; Deng et al. 2020), though it may form multimeric complexes with its closest homolog, MSL9 (Haswell et al. 2008)

Plants constitutively overexpressing *MSL10-GFP* display severe growth retardation, hyperaccumulation of H_2_O_2_, induction of genes associated with stress-response and cell death, and ectopic cell death (Veley et al. 2014; Basu et al. 2020). These phenotypes are reproduced in plants harboring an EMS-induced gain-of-function point mutation in the soluble C-terminus (S640L; *msl10-3G*) and in transgenic lines expressing an untagged version of *MSL10* containing seven Ser/Thr-to-Ala mutations in the soluble N-terminus from the native (or genomic) context (*MSL10g^7A^*) (Basu et al. 2020). Plants expressing wild-type *MSL10 (MSL10g)* or a phosphomimetic version of *MSL10* (*MSL10g^7D^*), do not share these phenotypes, suggesting that dephosphorylation of N-terminal Ser/Thr residues leads to channel activation (Basu et al. 2020). Because MSL10-GFP overexpressors, *msl10-3G* mutants, and plants expressing *MSL10g^7A^* all show the same gain-of-function phenotypes, we collectively refer to them here as *MSL10* hypermorphs. While the molecular mechanism by which *MSL10* hypermorphs are dysregulated has not yet been established, we have speculated that phosphorylation of the seven Ser/Thr residues in the MSL10 N-terminus keeps it in an inactive signaling state, and that either preventing phosphorylation with Ser/Thr-to-Ala mutations or overexpression of the GFP-tagged protein circumvents this regulation (Veley et al. 2014; Basu et al. 2020). We also recently proposed that an interaction between the soluble N- and C-termini of MSL10 monomers accompanies MSL10 activation, and that the *msl10-3G* point mutation promotes or recapitulates this activation (Basu et al. 2020).

Dwarfism and spontaneous lesions are also observed in two previously characterized, partially overlapping, classes of *Arabidopsis thaliana* mutants. Autoimmune mutants are unable to properly suppress immune responses in the absence of pathogens (reviewed in (Wersch et al. 2016), while lesion mimic mutants exhibit uncontrolled programmed cell death on the leaf surface (reviewed in (Bruggeman et al. 2015)). Prompted by these similarities between autoimmune and lesion mimic mutants and *MSL10* hypermorphs, our previous characterizations of MSL10 as a sensor of membrane tension, and by the intriguing possibility that plants could sense mechanical signals of pathogen invasion, we investigated the potential role of MSL10 in plant defense responses against *Pseudomonas syringae*.

## RESULTS

### Growth at high temperatures suppresses the phenotypes associated with overexpression of *MSL10-GFP*, expression of *MSL10g^7A^*, and the *msl10-3G* mutant

Growth at elevated temperatures (28-30°C) can suppress the dwarfing and the localized ectopic cell death in autoimmune mutants (Zhu et al. 2010; Huot et al. 2017; Hammoudi et al. 2018), and we speculated that the same might be true for *MSL10* hypermorphs. As shown previously (Veley et al. 2014), MSL10-GFP overexpressors were dramatically smaller than the WT when grown at 21°C (**Figure 1A, B**). However, when grown at high temperature, *MSL10-GFP* overexpressing lines were nearly indistinguishable from wild-type, *msl10-1*, or *msl9-1 msl10-1* mutant plants in terms of rosette size and fresh weight. Elevated temperature did not substantially alter the level of transcripts from the endogenous *MSL10* gene in wild-type plants, *msl10-1* null mutants, or *MSL10-GFP* overexpression lines (**Figure 1C**), and MSL10-GFP protein abundance was unaffected (**Figure 1D**). Furthermore, MSL10-GFP was localized at the periphery of the cell at both normal and high temperatures in all three transgenic lines (**Figure 1E**). Growth at high temperature also suppressed dwarfing in *msl10-3G* mutants and in plants expressing the *MSL10g^7A^* transgene but did not affect the growth of plants expressing *MSL10g* or *MSL10g^7D^*. High temperature did not appreciably affect transcript levels of endogenous or transgenic *MSL10* in these lines (**Figure S1**). Thus, growth at elevated temperatures relieved dwarfing in all three *MSL10* hypermorphs but did not do so by altering transcript levels, protein abundance or subcellular localization. Growth at elevated temperature also suppressed previously documented cellular phenotypes of *MSL10* hypermorphs, including the hyperaccumulation of reactive oxygen species, ectopic cell death, and increased transcript levels of four genes previously established as hallmarks of MSL10 activity (Veley et al. 2014). We assessed these phenotypes in rosette leaves from three-week-old wild-type, *msl10-1* and *msl10-3G* mutant plants; three independent transgenic lines overexpressing *MSL10-GFP* (previously described in (Veley et al. 2014)); and *msl10-1* plants harboring *MSL10g, MSL10g^7A^* or *MSL10g^7D^* transgenes (which contain a genomic copy of wild-type *MSL10, MSL10* with seven phospho-mimetic mutations in the soluble N-terminus, or *MSL10* with seven phospho-dead mutations in the soluble N-terminus, respectively, all previously described in (Basu et al. 2020)). When plants were grown at high temperatures, the increased O^2-^, H_2_O_2_, and ectopic cell death observed in *MSL10-GFP* overexpression lines, the transgenic lines were similar to that found in the wild type (**Figure 2A-C**). *MSL10g* and *MSL10g^7D^* lines and *msl10-1* mutants were indistinguishable from the wild-type when grown at high or at normal temperatures. The difference in scale between the data shown in the two panels of Figure 2B and C can be attributed to different instruments used in analyzing ROS and cell death (See Methods). Finally, the transcript levels of four genes previously identified as reporters of *MSL10* over-activation (Veley et al. 2014; Basu et al. 2020) were reduced to almost wild-type levels when *MSL10-GFP* overexpression plants were grown at high temperatures (**Figure 2D**). Taken together, these data show that growth at high temperature relieves all the phenotypes associated with the presence of hypermorphic alleles of *MSL10*, a characteristic shared with autoimmune mutants.

**Figure 1.**
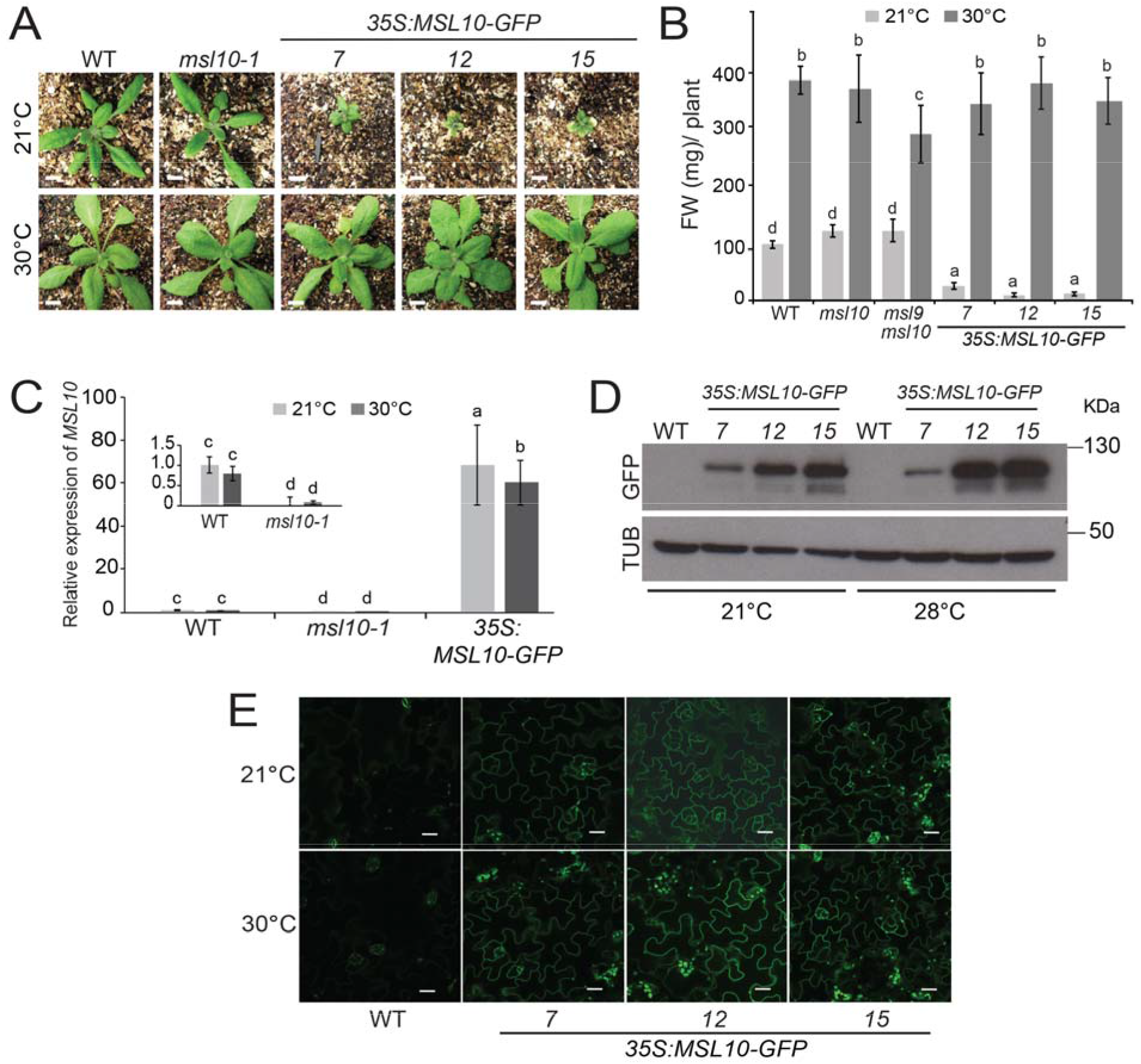
Growth at high temperatures suppresses dwarfing in *MSL10-GFP* overexpressors. **(A)** Three-week-old seedlings from wild-type, *msl10-1*, and three independent T4 homozygous lines overexpressing MSL10-GFP, grown side-by-side on soil under 24 h of light at 21°C or 30°C. Bars, 0.5 cm. **(B)** Average fresh weight (FW) of the aerial portion of plants grown as in (A). Error bars indicate SD of three independent trials, 10 leaves per trial. **(C)** *MSL10* transcript abundance relative to the wild type in wild-type, *msl10-1*, or *35S:MSL10-GFP-15* plants, grown as in (A). Inset, wild-type and *msl10-1* lines only. Expression levels were normalized to *EF1a* and *UBQ5*. Error bars indicate SEM from three independent trials, three technical replicates per trial. **(D)** Immunoblot of protein extracts from rosette leaves of two-week-old plants. MSL10-GFP was detected with an anti-GFP primary antibody (top), then the blot was stripped and re-probed with an anti-α tubulin primary antibody (bottom). **(E)** Confocal images of leaf epidermal cells from plants grown as in (A). GFP signal is shown in green. All images were obtained using identical settings. Bars, 20 μm. In B and C, different letters indicate significant differences as determined by two-way ANOVA followed by Tukey’s post-hoc test (P < 0.05).

**Figure 2.**
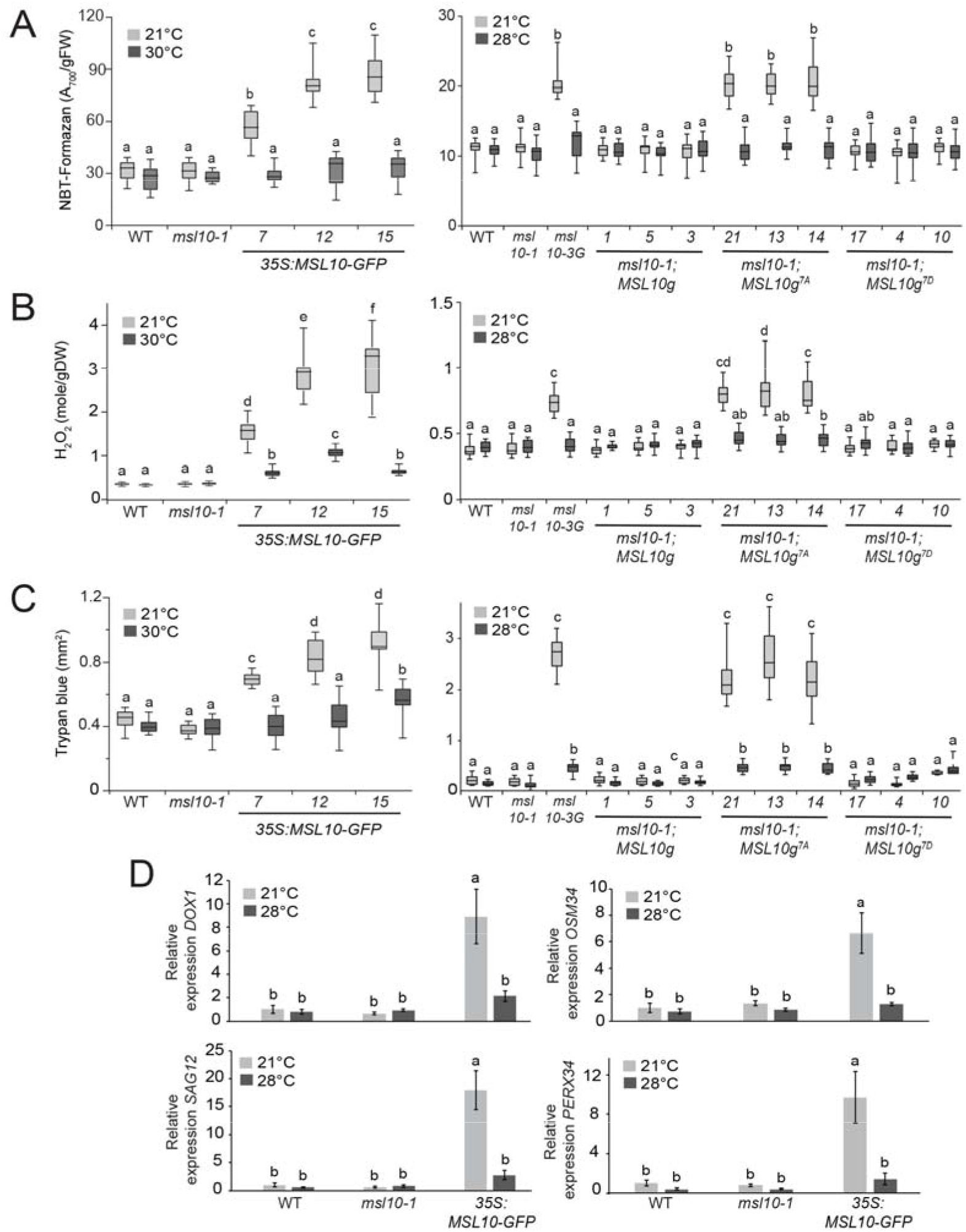
Growth at high temperatures suppresses multiple cellular and molecular phenotypes associated with *MSL10* hypermorphs. **(A)** Colorimetric quantification of superoxide content, as indicated by NBT-formazan deposition, in rosette leaves in three-week-old plants of the indicated genotypes grown at 21°C or 28° C under 24 hours of light. **(B)** Hydrogen peroxide (H_2_O_2_) content in rosette leaves grown as in (A), quantified with the Amplex Red-coupled quantitative assay. **(C)** Ectopic cell death indicated by Trypan blue staining in rosette leaves grown as in (A). (**D**) Relative transcript abundance of four previously characterized stress-responsive genes. Plant growth, cDNA synthesis, quantification, and statistical analysis were performed as in Figure 1C. In all panels, different letters indicate significant differences as determined by two-way ANOVA followed by Tukey’s post-hoc test (P < 0.05). A-C present the means ± SD from three independent trials, 5-10 leaves per genotype.

### *MSL10-GFP* Overexpression Phenotypes Require *SGT1b, RAR1*, and *HSP90.1*

Most NLRs require *SUPPRESSOR of G2 ALLELE of SKP1* (*SGT1*) for function (Kadota et al. 2010). SGT1 is part of a co-chaperone complex that includes REQUIRED for MLAI2 RESISTANCE (RAR1), and HEAT SHOCK PROTEIN 90 kD (HSP90) (Siligardi et al. 2018; Takahashi et al. 2003; Liu et al. 2004). SGT, RAR1 and HSP90 interact directly with NLRs and are required for their accumulation and subcellular trafficking (reviewed in (Borrelli et al. 2018)). Thus, in the absence of any known NLR associated with MSL10-induced downstream autoimmune response or any evidence of MSL10 acting as a target for a pathogen effector protein, we tested if silencing of *SGT1, RAR1*, and *HSP90* by <u>V</u>irus <u>I</u>nduced <u>G</u>ene <u>S</u>ilencing or VIGS (Baulcombe 1999) could abolish the cell death triggered by transiently overexpressed *MSL10-GFP* in *Nicotiana benthamiana*.

Leaves were agroinfiltrated with VIGS constructs (*TRV2:SGT1, TRV2:RAR1, TRV2:HSP90*, or empty vector). After ten days, transcript levels of each target gene were reduced by ~80% (**Figure S2A**). After another ten days, leaves were subjected to a second agroinfiltration with *35S:MSL10-GFP, 35S:MSL10^7D^-GFP*, or *35S:MSL10^7A^-GFP*. Cell death was quantified 5 days later, using dual FDA/PI staining as described (Veley et al. 2014). As expected, in leaves that had been pre-infiltrated with an empty VIGS vector, transient expression of wild-type *MSL10-GFP* resulted in the death of ~27% of *N. benthamiana* epidermal cells, and expression of *MSL10^7A^-GFP* triggered even more cell death (~39%), while expression of *MSL10^7D^-GFP* resulted in the death of just 11% of cells (**Figure 3A**). However, we found that silencing of either *SGT1, RAR1*, or *HSP90* almost completely prevented cell death in leaves that were transiently expressing *MSL10-GFP* (~10%) or *MSL10^7A^-GFP* (~16%). Immunoblotting showed that these differences in cell death could not be attributed to differences in MSL10-GFP protein abundance (**Figure S2B-D**).

We next investigated the same question in Arabidopsis plants stably expressing *35S:MSL10-GFP* into the T-DNA insertion lines *sgt1b-1* (SALK_026606) and *hsp90.1-1* (SALK_007614; (Takahashi et al. 2003) and the point mutant *rar1-21* (Tornero et al. 2002)(Tornero et al., 2002)(Tornero et al., 2002). The *sgt1b-1* mutant line has a T-DNA inserted in the promoter region of the *SGT1b* gene and *SGT1b* expression was reduced to approximately 60% of that in wild-type plants (**Figure S3A-B**), indicating that the *sgt1b-1* mutant is a knock-down allele of *SGT1b*. The *sgt1b-1, rar1-21*, and *hsp90.1* mutants did not exhibit detectable morphological differences compared to wild-type plants (**Figure S3C**). To avoid variability in transgene expression due to position effect, we decided to first incorporate a transgene in a mutant background and cross the T1 selected plant with wild-type Col-0. The segregating F2 progenies of homozygous knock-out mutant and wild-type (both harboring the transgene integrated in the same location in the chromosome of either genetic background) were considered for further analysis.

**Figure 3.**
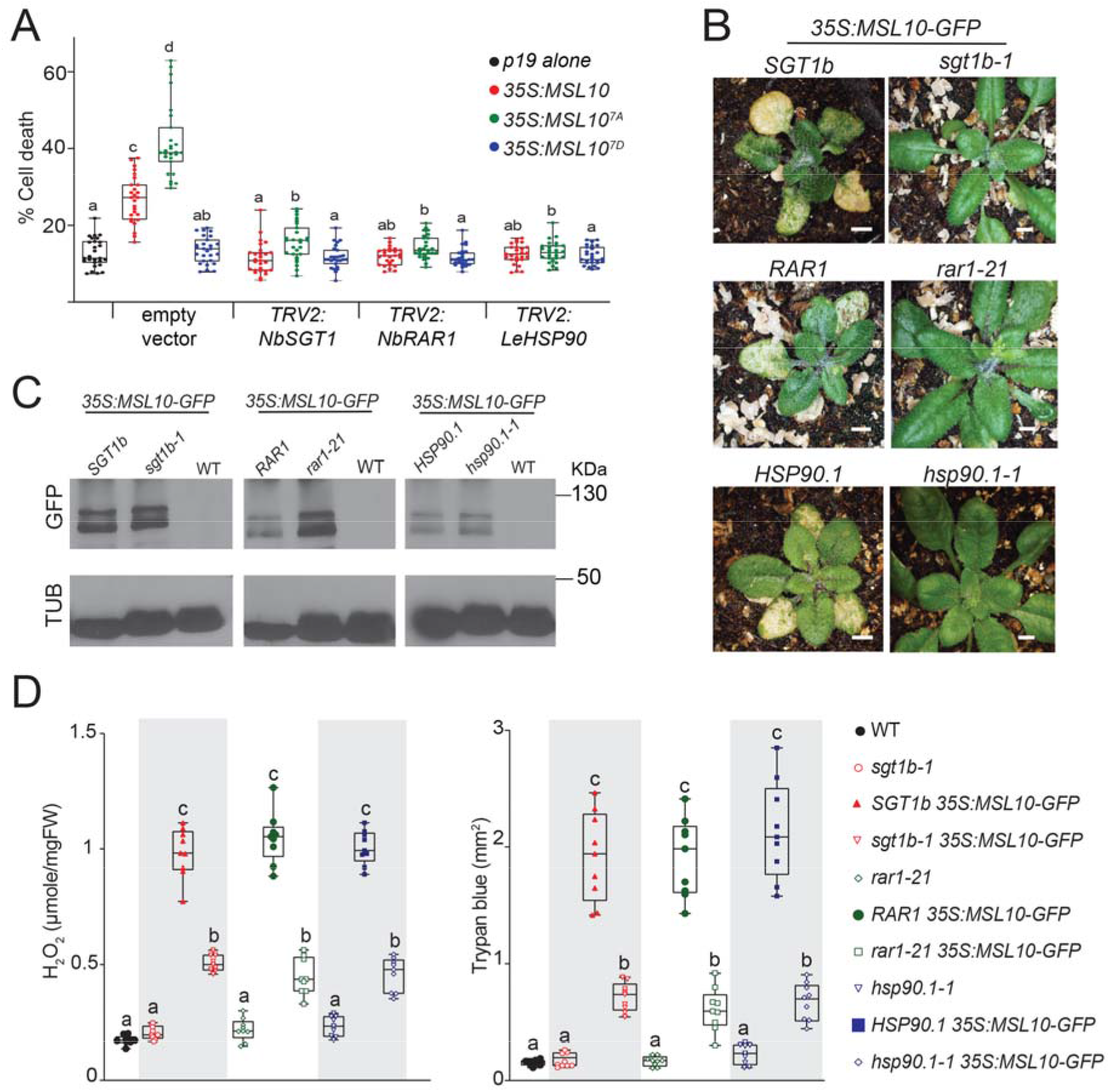
*SGT1b, RAR1* and *HSP90* are required for phenotypes associated with *MSL10-GFP* overexpression in tobacco and Arabidopsis. **(A)** Quantification of cell death, as determined by dual staining of FDA and PI, in epidermal cells from *N. benthamiana* leaves transiently overexpressing MSL10-GFP and its phospho-variants when *SGT1b, RAR1*, or *HSP90* was silenced. The means and SD from three independent trials, ~300 epidermal cells per trial, are presented. **(B)** *35S:MSL10-GFP* seedlings segregating *sgt1b-1, rar1-21*, or *hsp90.1-1*, grown side-by-side on soil under 24 h light at 21°C. Left panel bar, 0.5 cm; right panel bar, 0.35 cm. **(C)** Immunoblot showing MSL10-GFP levels in total protein extracted from three-week-old rosette leaves grown as in (B). **(D)** Hydrogen peroxide (left) and Trypan blue (right) content in rosette leaves, quantified as in Figure 2. The means and SD from three independent trials, each with 3 leaves are presented. In A and D, different letters indicate significant differences as determined by two-way ANOVA followed by Tukey’s post-hoc test (P < 0.05).

As expected, plants overexpressing *35S:MSL10-GFP* in the wild type *SGT1b, RAR1*, and *HSP90.1* backgrounds exhibited dwarfing and ectopic cell death as assessed by the appearance of yellowish-brown lesions on rosette leaves. (**Figure 3B**). However, these phenotypes were largely absent in *MSL10-GFP* overexpressing siblings homozygous for *sgt1b-1, rar1-21*, and *hsp90.1-1*. Immunoblotting and confocal imaging showed that neither MSL10-GFP protein abundance nor localization were altered in these backgrounds (**Figure 3C, S3D**). H_2_O_2_ hyperaccumulation and ectopic cell death were also markedly reduced in *sgt1b-1; 35S:MSL10-GFP* (~50% and 65%), *rar1-21; 35S:MSL10-GFP* (~60% and 70%), and *hsp90.1-1; 35S:MSL10-GFP* (~50% and 67%) lines compared to their *35S:MSL10-GFP* siblings (**Figure 3D**). We obtained similar results with a recently described null allele of *SGT1b* (*sgt1b-2*, GABI_857A04, (Zhang et al. 2015) (**Figure S3E-G**). These results collectively indicate that the dwarfing, H_2_O_2_ accumulation and ectopic cell death associated with *MSL10-GFP* overexpression require the surveillance complex SGT1/RAR1/HSP90.1 in both *N. benthamiana* and Arabidopsis. However, this effect does not appear to be mediated through protein-protein interactions, as SGT1b, RAR1, and HSP90.1 did not associate with MSL10 in bimolecular fluorescence complementation assays (**Figure S4**).

### The phenotypes associated with MSL10-GFP overexpression are independent of *EDS1* and *NDR1*

Downstream of NLR function, the immune regulator ENHANCED DISEASE SUSCEPTIBILITY (EDS1) is a lipase-like protein that serves to connect TIR-NLR signaling with downstream events including programmed cell death, production of the defense hormone salicylic acid (SA), and defense gene expression (reviewed in (Dongus and Parker 2021)). NONRACE-SPECIFIC DISEASE RESISTANCE 1 (NDR1), a membrane-associated integrin-like protein, plays a similar role for CC-NLRs (Century et al. 1997; Knepper et al. 2011; Coppinger et al. 2004; Century et al. 1995). To investigate whether disruption of *EDS1* or *NDR1* are required for dwarfing or cell death observed in MSL10-GFP overexpression lines or the *msl10-3G* background, we introduced *35S:MSL10-GFP* into the *eds1-23* (SALK_057149, (Song 2016)) and the *ndr1-1* (Century et al. 1997) null mutant backgrounds by transformation. As anticipated, expression of *35S:MSL10-GFP* in wild-type plants resulted in severe dwarfing and the appearance of yellowish-brown lesions on rosette leaves (**Figure 4A, top panels**), and Trypan blue-stained patches on leaves (**Figure 4A, bottom panels**) compared to the wild type or to untransformed *eds1-23* or *ndr1-1* mutants. We found that neither the *eds1-23* nor the *ndr1-1* mutant background appreciably affected the phenotypic results of *MSL10-GFP* overexpression, and immunoblotting confirmed similar MSL10-GFP levels in all lines (**Figure 4B**). Similarly, neither the *eds1-23* nor the *ndr1-1* alleles altered dwarfing or ectopic cell death observed in the *msl10-3G* background (**Figure 4C**). Collectively, these results indicate that the phenotypes associated with *MSL10* hypermorphs are independent of *EDS1* and *NDR1*.

**Figure 4.**
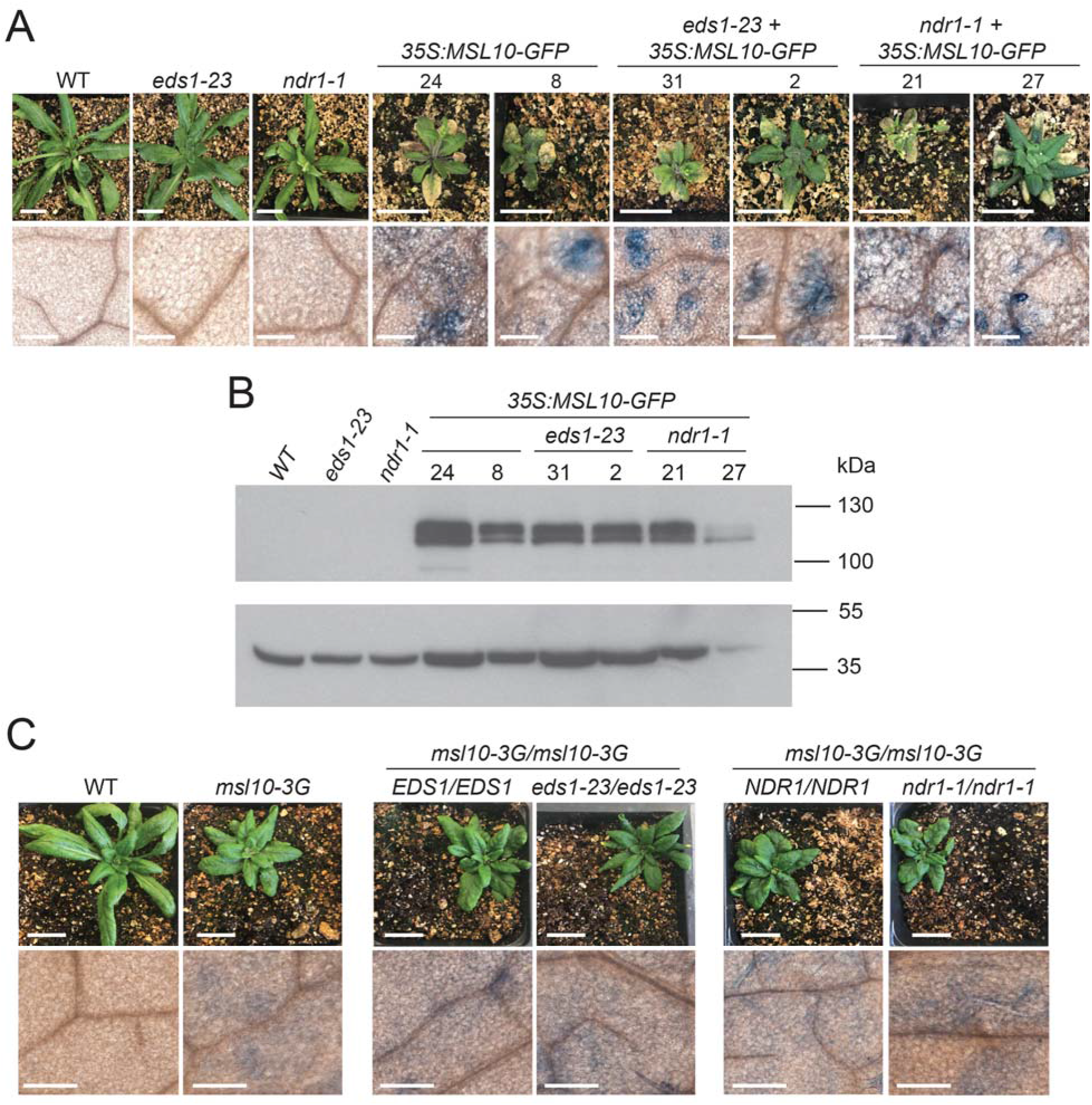
Disruption of *EDS1* or *NDR1* fails to suppress phenotypes associated with *MSL10* hypermorphs. **(A)** Top, five-week-old T1 plants grown side-by-side in 24 h of light at 21°C. Bars, 2 cm. Bottom, rosette leaves from the same plants stained with Trypan blue to visualize dead cells. Bars, 200 μm. **(B)** Immunoblot of protein extracts from leaf tissue of T1 lines, performed as in Figure 1D. **(C)** *msl10-3G* siblings segregating *eds1-23* or *ndr1-1* alleles. Top, four-week-old plants grown as in (A). Bars, 2 cm. Bottom, dead cells in rosette leaves visualized by Trypan blue staining. Bars, 200 μm.

### *PAD4* and *SID2* are partially required for the dwarfing and ectopic cell death associated with MSL10-GFP overexpression

SA accumulation is often associated with inhibition of growth, spontaneous cell death, and immune responses (Bruggeman et al. 2015; Peng et al. 2021). This prompted us to investigate whether SA biosynthesis might be required for MSL10-associated phenotypes. Towards this goal, we crossed plants expressing *35S:MSL10-GFP* with *PHYTOALEXIN DEFICIENT 4-1 (pad4-1), SA INDUCTION DEFICIENT2-2 (sid2-2)* double mutants (Jirage et al. 1999; Wildermuth et al. 2001). *SID2* gene encodes ICS1, a key SA biosynthetic enzyme, while PAD4 is a lipase-like proteins that is involved transcriptional upregulation of SA accumulation (Ng et al. 2011). We followed the approach described above to assess transgenic siblings with different genetic backgrounds. Four-week-old *pad4-1sid2-2* siblings were phenotypically indistinguishable from the wild-type parental line, but *pad4-1sid2-2* siblings overexpressing *MSL10-GFP* displayed partial suppression of the dwarfing and fresh weight seen in the *35S:MSL10-GFP* line (**Figure 5A-B**). Consistent with these phenotypic features, the level of ectopic cell death in *pad4-1sid2-2; 35S:MSL10-GFP* leaves was ~ 50 % lower than that observed in *35S:MSL10-GFP* overexpression lines (**Figure 5C**). MSL10-GFP levels were comparable in both types of siblings as assessed by immunoblotting (**Figure 5D**). These results suggest that SA biosynthesis influences but is not required for the autoimmune phenotypes associated with *MSL10-GFP* overexpression.

**Figure 5.**
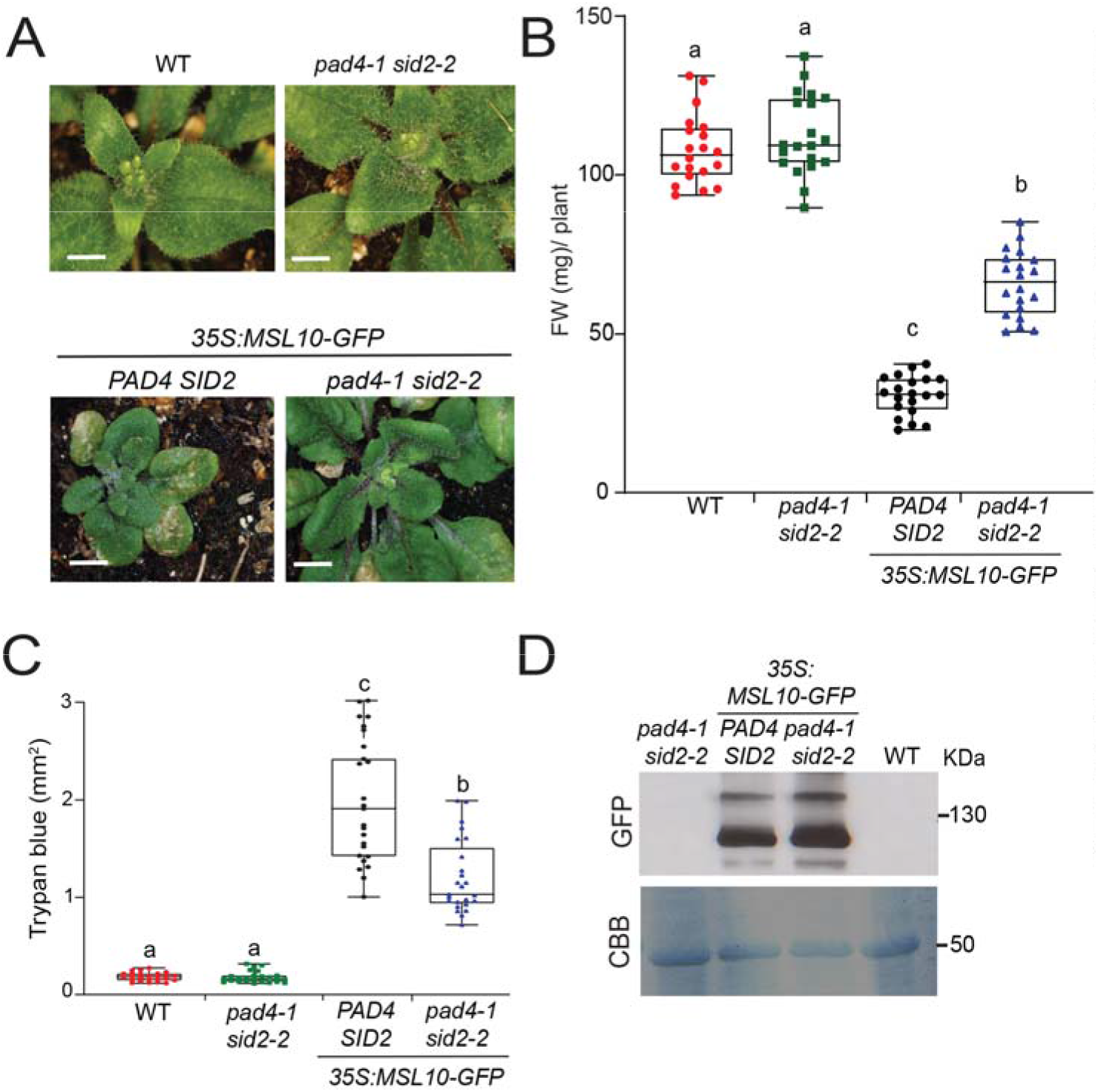
*PAD4* and *SID2* are partially required for the phenotypes associated with overexpression of *MSL10-GFP*. **(A)** Four-week-old seedlings grown side-by-side in 24 h of light at 21°C. Bar, 1 cm. **(B)** Average fresh weight (FW) of rosettes grown as in (A). **(C)** Ectopic cell death indicated by Trypan blue staining in rosette leaves. **(D)** Immunoblot of MSL10-GFP protein levels from three-week old rosette leaves grown as in A, performed as in Figure 1D. In B and C, the mean and SD of three independent trials, 8-10 leaves each are presented, and different letters indicate significant differences as determined by two-way ANOVA followed by Tukey’s post-hoc test (P < 0.05).

### Hypermorphic alleles of *MSL10* confer reduced susceptibility to *Pseudomonas syringae*

*Pseudomonas syringae* is a model for plant pathogen studies, and programmed cell death is commonly triggered during the hypersensitive response of Arabidopsis to *Pseudomonas* syringae interactions (Xin and He 2013). Since MSL10 hypermorphs are known to trigger cell death in response to cell swelling (Basu and Haswell 2020), and cell death is a typical immune response, we asked whether *MSL10* contributes to pathogen resistance. Plants were grown at 28°C under short-day conditions for 5-6 weeks to suppress the cell death and dwarfing phenotypes of *MSL10* hypermorphs prior to infection. Then, since elevated temperature suppresses both plant immunity and pathogen virulence (Huot et al. 2017), 5-to 6-week-old plants were transferred to 21 °C twenty-four hours before inoculation with bacterial cultures. Using this strategy, as shown in Figure S5, the levels of H_2_O_2_ and cell death plants harboring *MSL10* hypermorphic alleles remained indistinguishable from the wild-type plants at the time of infection, 1 day after transfer to 21°C.

Fully expanded leaves were syringe-inoculated with *Pto* DC3000, which is virulent and can multiply to high levels (100 to 1000-fold) in wild-type Col-0 plants. At the start of the experiment, no differences in bacterial counts were observed among any lines (**Figure 6A**). At 3 days post-infection, *msl10-3G* mutants supported significantly lower bacterial growth than wildtype plants, as did plants overexpressing *MSL10-GFP* and plants expressing *MSL10g^7A^*. **In** contrast, phospho-mimetic versions of *MSL10* transgenes (*35S:MSL10-GFP^4D^* or *MSL10g^7D^*), and the *msl10-1* mutant had no apparent effect relative to the wild-type. The *pad4-1sid2-2* mutant, which does not accumulate SA in response to infection, exhibited enhanced susceptibility to *Pto* DC3000.

**Figure 6.**
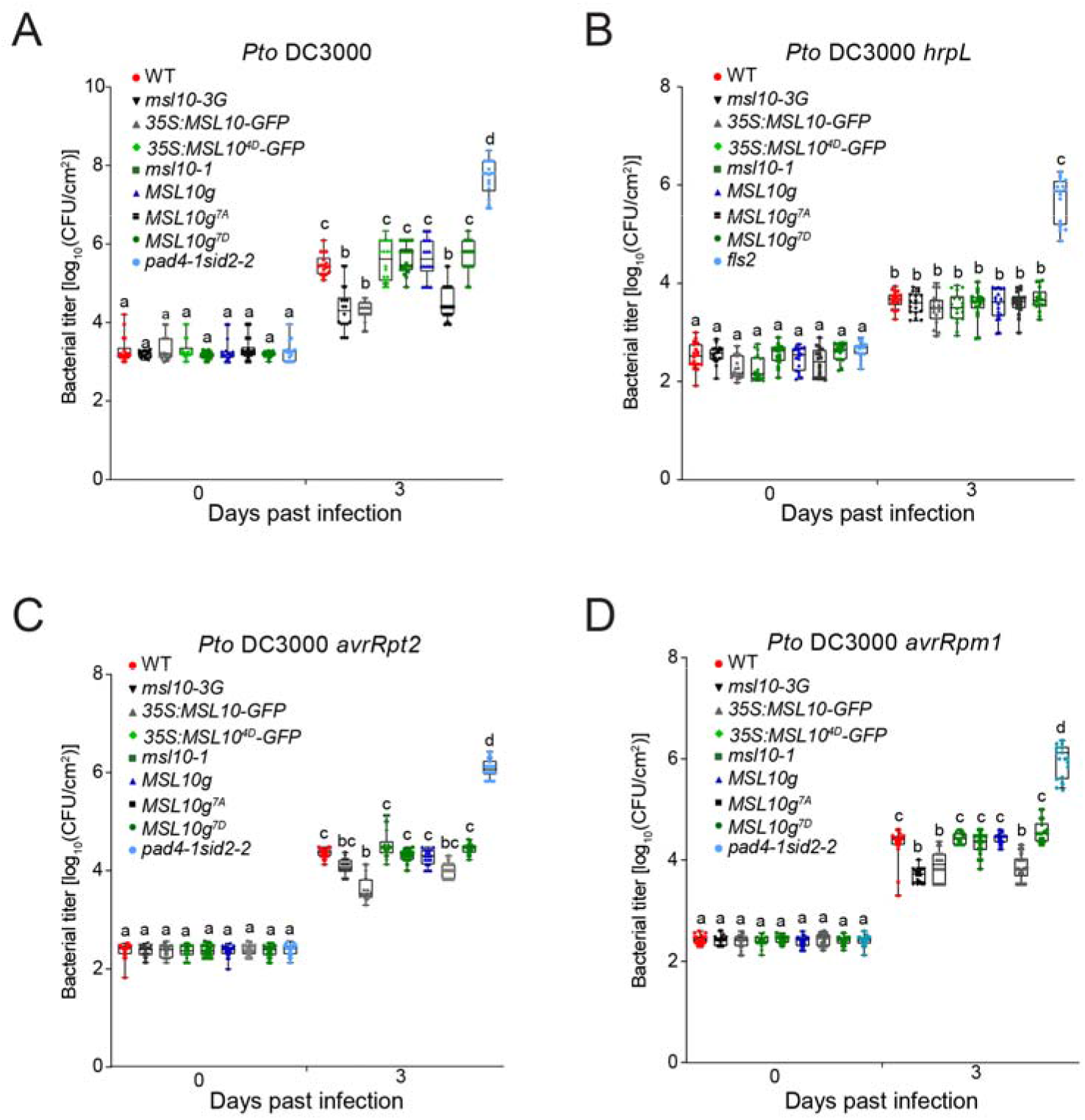
*MSL10* hypomorphs confer reduced susceptibility to *Pseudomonas syringae*. Bacterial growth in leaves from plants grown for 5 weeks side-by-side on soil at 28°C under a 10 h light /14 h dark regime, then transferred to 21°C for 1 day before syringe inoculation. Means and SD from three independent trials, each with 6 leaf extracts containing two leaf discs, per genotype, are presented. Different letters indicate a significant difference as determined by twoway ANOVA followed by Tukey’s post-hoc test (P < 0.05).

The enhanced resistance against *Pto* DC3000 exhibited in lines expressing *MSL10* hypermorphs suggested that MSL10 may be involved in basal plant defense responses. To test this idea, we syringe-inoculated plants with a mutant strain of *Pto* C3000, *hrpL*. This mutant strain lacks the HrpL transcription factor that controls the expression of type III secretion system genes required for effector secretion, cannot suppress basal defense, and therefore do not grow to high levels in wild-type Col-0 plants (Zwiesler Vollick et al. 2002). When inoculated with *Pto* DC3000 *hrpL* mutant bacteria, none of the lines showed apparent differences in viable bacterial counts (**Figure 6B**). In addition, we tested the resistance of these strains to avirulent *Pseudomonas syringae* expressing two well-characterized effectors, *Pto DC3000 avrRpt2* and *Pto DC3000 avrRpm1* (Mackey et al. 2003, 2002). We found that *MSL10* hypermorphs exhibited enhanced resistance to *Pto avrRpt2* (**Figure 6C**) or *Pto avrRpm1* (**Figure 6D**) compared to wildtype plants. However, *msl10-1* mutants and *MSL10g^7D^* lines were indistinguishable from the wild type in these assays. In summary, overactive *MSL10* alleles conferred reduced susceptibility to both virulent and avirulent strains of *P. syringae*, while *msl10-1* mutants and *MSL10g* lines were indistinguishable from the wild type in these assays. None of the lines tested impacted growth of the *hrpL* mutant. Taken together, these data suggest that *MSL10* hypermorphs may act as an autoimmune mutant.

### *MSL10* is required for normal *PR1* induction in response to *Pseudomonas* infection

The ectopic cell death observed in autoimmune mutants is often accompanied by constitutive upregulation of defense related genes like *PATHOGENESIS-RELATED 1* (*PR1*)(e.g., see (Li et al. 2010; Xu et al. 2015; Wang et al. 2017)). To investigate whether the same is true of *MSL10* hypermorphs, a time-course experiment was performed to evaluate *PR1* transcript accumulation in response to *Pto* DC3000 infection. We found that basal levels of *PR1* transcripts were the same in all lines tested. However, the kinetics of induction was different. In wild-type plants, increased levels of *PR1* transcripts were detected by 6 hours and continued to increase at 12- and 24-hour time points (**Figure 7A**). *PR1* transcript levels also increased after infection in *msl10-1* mutants, but the increase was not observable until 12 hours after treatment. On the other hand, *PR1* expression in the *msl10-3G* mutant (~18 to 45-fold) and in *MSL10-GFP* overexpression line (~24 to 57-fold), was higher than the wild type (~5 to 15-fold) as early as 6 h after *Pto* DC3000 infection. At all time-points tested, plants expressing phospho-mimetic *35S:MSL10^4D^-GFP* and *MSL10g* complemented lines displayed similar *PR1* expression kinetics as wild-type plants. We did observe small increases in *PR1* expression at later time points in all genotypes in response to mock infection. This may be attributed to mechanical or osmotic stress resulting from syringe-infiltration (Chandrashekar et al. 2018). Thus, MSL10 positively influences *PR1* induction after *Pto* DC3000 infection.

**Figure 7.**
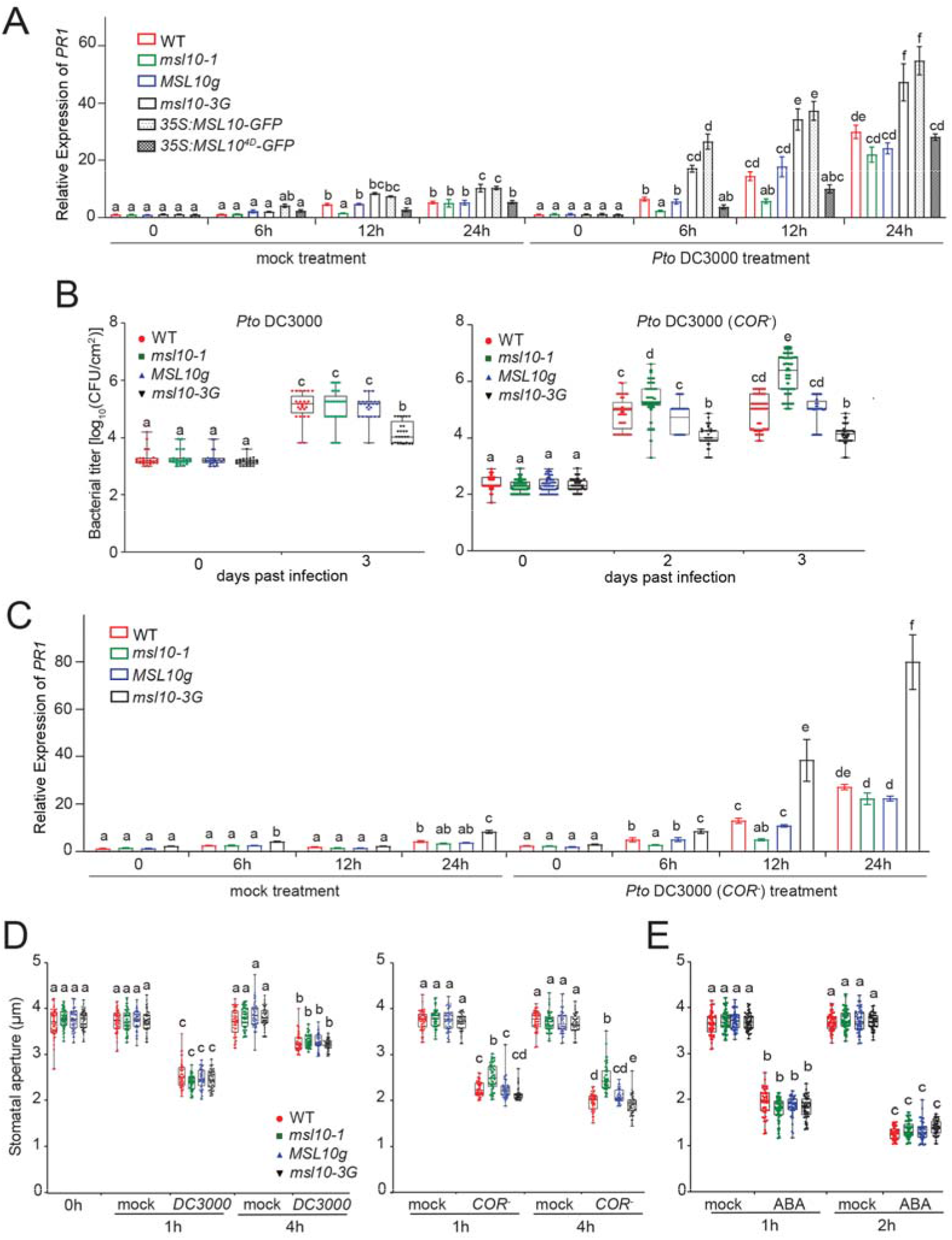
*MSL10* modulates the normal induction of *PR1*, susceptibility to *Pto* DC3000 (*COR^-^*), and stomatal closure. **(A)** *PR1* transcript abundance relative to the wild type, performed as in Figure 1C. RNA was isolated from plants grown as in Figure 6, then syringe inoculated with *Pto* DC3000. (**B**) Bacterial growth in leaves after spray inoculation with *Pto* DC3000 (*COR^-^*). The means and SD from five independent trials, each with 5-9 leaf extracts, are shown. Each extract contained two leaf discs. **(C)** *PR1* transcript abundance relative to the wild type, performed as in (A) except that plants were spray inoculated with *Pto* DC3000 (*COR^-^*). **(D, E)** Stomatal aperture measurements. Fully expanded, healthy leaves from 5-week-old plants grown as in Figure 6 were immersed in a suspension of *Pto* DC3000 or *Pto* DC3000 (*COR^-^*), or 10 μM ABA and stomatal apertures measured at the indicated time-points. Means and SD from three independent trials, each with 16 apertures, are shown. Different letters indicate a significant difference as determined by two-way ANOVA followed by Tukey’s post-hoc test (P < 0.05). CFU, colony-forming units; mock, 10 mM MgCl_2_.

### *MSL10* is required for normal basal defense, rapid *PR1* induction, and full stomatal closure in response to *Pto* DC3000 (COR^-^)

The observation that *msl10-1* mutants showed a delayed and attenuated induction of *PR1* led us to investigate the role of *MSL10* in responding to *Pto* infection under sensitized conditions. Under natural conditions, *Pseudomonas* enters plants either through wounds or natural openings like stomata and hydathodes (Melotto et al. 2008)). Syringe-infiltration directly delivers the bacteria into the apoplast, by-passing these natural entry mechanisms. To assess bacterial growth under more realistic conditions, we spray-inoculated plants with *Pto* DC3000. As with syringeinfiltration (**Figure 6A**), *msl10-3G* mutants displayed heightened resistance to *Pto* DC3000 compared to wild-type plants, while *msl10-1* mutants and the *MSL10g* line were indistinguishable from the wild-type. Coronatine deficient bacterial mutants (COR^-^) have severely compromised virulence when inoculated onto the leaf surface and are useful to characterize mutants with slightly weakened defense responses (Geng et al. 2014). When plants were spray-inoculated with *Pto* DC3000 (COR^-^) (Brooks et al. 2004), *msl10-3G* mutants again displayed enhanced resistance compared to the wild type at 2 or 3 days after infection (**Figure 7B**). Importantly, when using this weaker pathogen and spray inoculation, *msl10-1* showed reduced resistance compared to wild-type plants at both time points. We also observed a similar profile of *PR1* induction in response to infection with the *Pto* DC3000 (COR^-^) strain. *MSL10* hypermorphs exhibited higher levels of *PR1* transcripts, and *msl10-1* mutants displayed a delayed and reduced induction of *PR1* transcripts (**Figure 7C**). These results indicate that *MSL10* is required for normal levels of basal defense or innate immunity when presented with modest pathogenic challenges.

As *Pto COR^-^* strains are unable to inhibit stomatal closure and early PTI-mediated basal defense responses (Melotto et al. 2006), we hypothesized that the increased susceptibility in *msl10-1* mutants might be related to defects in stomatal closure. To test this hypothesis, we measured stomatal aperture from the leaf epidermis of wild-type, *msl10-1, MSL10g*, and *msl10-3G* lines exposed to *Pto* DC3000 (COR^-^). Wild-type leaves sprayed with *Pto* DC3000 briefly closed, then reopened, their stomates (**Figure 7D, left**), while the stomates of wild-type leaves spray-inoculated with *Pto* DC3000 (COR^-^) closed and did not open again (**Figure 7D, right**). All genotypes tested behaved similarly when spray inoculated with *Pto* DC3000. However, in response to *Pto* DC3000 (COR^-^), the stomates of *msl10-1* mutant were significantly more open than all the other genotypes both at 1 and at 4 hours after infection. ABA treatment efficiently produced stomatal closing in all genotypes after 2h of ABA incubation, further establishing that there is no generic defect in the ability of *msl10-1* stomates to close (**Figure 7E**).

## DISCUSSION

Considerable progress has been made in identifying and characterizing plant receptors involved in detecting bacterial effectors and MAMPs, but less is known about non-canonical players involved in plant defense. Here we investigated the role of the mechanosensitive ion channel MSL10 in plant immunity. We found that growth at elevated temperature, loss of the SGT1/RAR1/HSP90 chaperone complex, or disruption of SA biosynthesis fully or partially alleviated all the phenotypes triggered by *MSL10* hypermorphic alleles that we tested. These phenotypes, most of which are associated with enhanced basal defenses, included dwarfing, ectopic cell death, hyperaccumulation of ROS, and a characteristic gene expression profile (**Figure 1–3, 5, Figure S1-3**). However, the NLR signal transducers EDS1 and NDR1 were not required (**Figure 6**). *MSL10* hypermorphs exhibited enhanced disease resistance against both virulent and avirulent strains of *Pseudomonas syringe*, but not *hrpL* (**Figure 6**), and *msl10* null mutants showed increased susceptibility to infection by coronatine-deficient *Pto* (**Figure 7**). We also found that MSL10 promotes precocious *PR1* induction and is required for normal stomatal closure in response to *Pto* DC3000 (COR^-^) infection (**Figure 7**).

### MSL10 hypermorphs compared to lesion mimic and autoimmune mutants

*MSL10* hypermorphs strongly resemble a category of mutants called Lesion Mimic Mutants (LMMs). LLMs display spontaneous cell death in the absence of any pathogenic infection, mechanical stress, or abiotic stress (Moeder and Yoshioka 2008; Bruggeman et al. 2015). Like *MSL10* hypermorphs, LMMs often share other phenotypes, including dwarfing and the constitutive activation of defense responses. Also similar to *MSL10* hypermorphs, the phenotypes of some LMMs are temperature-sensitive (Huang et al. 2010; Yang and Hua 2004; Yang et al. 2010). Some LMMs are autoimmune mutants--that is, their phenotypes result from the loss of repression of pathogen responses (Wersch et al. 2016; Chakraborty et al. 2018). Autoimmune effects can result from the overactivation of PRRs and NLR proteins, the loss of negative regulators of NLRs, and/or the disruption of NLR-guardee interactions and mis-regulation of SA biosynthesis. In this study, we provide evidence that the LMM-like phenotypes exhibited by *MSL10* hypermorphs are due to the induction of autoimmune responses (see discussion below). However, unlike canonical/known autoimmune mutants, *MSL10* hypermorphs did not exhibit increased basal expression of the *PR1* gene (**Figure 7A**). Instead, they might more appropriately be considered constitutive priming mutants, poised to respond to the presence of a pathogen more quickly or to a higher degree than the wild type (Mauch-Mani et al. 2016)

### MSL10 and R gene-mediated immunity

The SGT1b/RAR1/HSP90 co-chaperone complex is known to function as an integral component of ETI (Wersch et al. 2019). Silencing of SGT1/RAR1/HSP90 components alleviates autoimmune responses in several autoimmune mutants. For example, SGT1b is a positive regulator of the autoimmune response displayed by *chs3* mutants (Xu et al. 2015), HSP90.3 is required for the constitutive activation of defense response in the *saul1-1* mutant (Liang et al. 2020) and *topp4-1* autoimmune phenotypes require RAR1 and HSP90.2 (Yan et al. 2019). Similarly, the spontaneous cell death and hyperaccumulation of ROS exhibited by plants overexpressing MSL10-GFP required *SGT1b, RAR1*, and *HSP90* in both tobacco and Arabidopsis (**Figure 3, S2** and **S3**).

As we were unable to detect a physical interaction between MSL10 and SGT1b, RAR1 or HSP90.1 (**Figure S4**), MSL10 is unlikely to be a client of the co-chaperone complex. Rather, these data suggest that MSL10 acts upstream of R gene-mediated immune responses. However, most NLR-mediated signaling requires either *EDS1* or *NDR1*, and we observed that disruption of *EDS1* and *NDR1* did not alleviate the autoimmune-like phenotypes associated with *MSL10* hypermorphs. Furthermore, while autoimmunity can be caused by defects in guardee proteins, *msl10* null mutants do not show an autoimmune response under unstressed conditions, indicating that MSL10 is unlikely to function as a guardee. A few scenarios fit the data presented here: 1) MSL10 could function upstream of a pathway that involves R proteins that are *EDS1-* and *NDR1*-independent. There is at least one other example of an immune response that requires *SGT1* but not *EDS1* (Sang et al. 2020); 2) the MSL10-induced autoimmunity pathway could involve non-NLR R proteins; or 3) MSL10 could function in a pathway that is entirely independent of R genes. For the latter to be the case, the SGT1 complex would have to be involved in an R-gene independent function as well, and both HSP90 and SGT1b have been shown to interact with the receptors for plant hormones auxin and jasmonic acid (Wang et al. 2016; Zhang et al. 2015).

### MS ion channels in pathogen perception

Cation fluxes are immediate and essential parts of the host immune response (Moeder et al. 2018), and it was recently reported that the OSCA1 cation channel is required for defense-induced stomatal closure (Thor et al. 2020). While MSL10 is unlikely to transport calcium directly (Maksaev and Haswell 2012), the release of anions through MSL10 could depolarize the plasma membrane, thereby indirectly activating calcium channels (Guerringue et al. 2018). Indeed, MSL10 is closely associated with and required for the cytoplasmic calcium transient associated with cell swelling (Basu and Haswell 2020). However, the ability to trigger dwarfing and programmed cell death may be triggered independently of ion flux through MSL10 (Veley et al. 2014; Maksaev et al. 2018). Thus, it is possible that MSL10 serves as a sensor of membrane integrity that is triggered by changes in membrane tension resulting from *Pseudomonas* infection, either due to the insertion/assembly of the type III secretion apparatus within the plasma membrane, or indirectly to events triggered by cytoplasmic effectors such as water-soaking (though water-soaking is not seen with avirulent strains (Xin et al. 2016)). Activation of MSL10 could then lead to calcium transients or to non-conducting interactions with components of R gene-mediated immunity, eventually leading to *PR1* induction and/or stomatal closure. Interestingly, the putative MS ion channel MSL4, a close homolog of MSL10, interacts with the defense regulator ACD6 and is required for PRR-mediated basal immunity responses (Zhang et al. 2017), and AtPIEZO1 is required for long distance virus translocation (Zhang et al. 2019). Taken together, these results suggest that MS ion channels play diverse, yet-to-be-discovered roles in plant defense.

## MATERIALS AND METHODS

### Plant lines and growth conditions

*Arabidopsis thaliana* plants used in this study were of the Col-0 ecotype. The following mutants were used: *sid2-2* (Wildermuth et al. 2001)(Wildermuth et al., 2001)(Wildermuth et al., 2001), *pad4-1* (Jirage et al. 1999), *eds1-23* (SALK_057149; (Song 2016), *rar1-21* (Tornero et al. 2002), *sgt1b-2* (GABI_857A04; (Zhang et al. 2015)), *sgt1b-1* (SALK_026606), *ndr1-1* (Century et al. 1997), (*hsp90.1-1* (SALK_007614; (Takahashi et al. 2003)), *fls2* (salk_141277; (Guo et al. 2014)), *msl10-1* (SALK_114626), *msl10-3G* (Zou et al. 2016; Basu et al. 2020), and *msl9-1* (SALK_114361; (Haswell et al. 2008)). In most experiments, plants were grown on soil at 21°C under a 24-h light regime (~120 μmol m^-2^s^-1^). For experiments done at elevated temperature, plants were grown on soil at high temperatures that ranged from 28°C to 30°C with 24-h light. For bacterial infection assays and stomatal aperture measurements, plants were grown at 21°C under a 24-h light regime for 10 days until the seedling germinated followed by 28°C under short-day (10 h of light/14 h of dark) for 5weeks. Plants were then acclimatized for 24-h at 21°C in short-day conditions before subjecting them to bacterial infection. The *MSL10-GFP* overexpression lines (line #7-1, # 12-3 and #15-2) and *MSL10g, MSL10g^7A^*, and *MSL10g^7D^* plant lines are described in earlier studies (Veley et al. 2014; Basu et al. 2020).

### Generation of transgenic lines

The *35S:MSL10-GFP* plasmid (Veley et al. 2014) was transformed into Col-0 (wild type), *sgt1b, rar1-21, hsp90.1-1, pad4-1sid2-2, eds1-23*, or *ndr1-1* mutant backgrounds by *Agrobacterium tumefaciens* (GV3101) mediated floral dipping (Clough and Bent 1998). Transgenic plants were selected on soil by spraying with 200 mg/L Basta followed by segregation analysis and selection of homozygous transgenic lines with single insertion of transgene on MS plates supplemented with 20 mg/L Basta. Sequences of the primers used for genotyping transgenes, or the mutant background are listed in **Supplemental Table S1**. Since transgene expression in a knock-out mutant background relative to the transgene expression in the wild-type plant does not involve identical genetic background, it is difficult to compare phenotypes among the transgenic plants. To eliminate such issues, we first introduced the transgene into the knock-out mutant line and then crossed the T1 transgenic line with the wild-type Col-0 line. Segregating homozygous F2 siblings with identical genetic backgrounds were used for comparison. Plants with the *msl10-3G* allele were crossed to *eds1-23* and *nrdr1-1* plants. F2 progeny homozygous for the *msl10-3G* allele were identified by genotyping, and comparisons were made between those F2 siblings homozygous for wild-type or mutant *eds1-23* and *nrdr1-1* alleles.

### Gene expression analysis

Quantitative reverse-transcription polymerase chain reaction (qRT-PCR) was performed as described (Basu et al. 2020) with minor modifications. In most experiments, total RNA was isolated from the rosette leaves (5^th^ and 9^th^) of healthy 3-week-old plants using the RNeasy Plant Mini Kit (Qiagen) including DNase I treatment. Rosette leaves from five-to-six-week-old plants were used for RNA extraction for transcript analysis in response to bacterial infection. Reverse transcription was performed using 2 μg of total RNA and oligo(dT)_20_ primers with M-MLV Reverse Transcriptase (Promega). All results shown include data from three independent experiments; for each treatment three technical replicates were performed for each of three independent experiments. Expression levels were normalized to the geometric mean of two reference genes, *UBQ5* and *EF1α*. Finally, the relative abundance of transcripts was calculated using the 2^−ΔΔ*Ct*^ method (Livak and Schmittgen 2001). Semi-quantitative RT-PCR was performed as in (Veley et al. 2014). Sequences of the primers used are listed in **Supplemental Table S1**.

### Quantification of ROS in tissue extracts

H_2_O_2_ levels in leaf extracts were measured using the Amplex Red Hydrogen Peroxide/Peroxidase Assay Kit (Molecular Probes, Invitrogen) (Basu et al. 2020). Three to five independent experiments were used for each condition at each time interval. The leaves were weighed, frozen and homogenized in phosphate buffer. Homogenates were centrifuged at 10,000 g for 10 min at 4°C and the supernatant was used in the Amplex red assay according to the manufacturer’s instructions. Samples were measured with a 96-well microplate reader (Infinite 200 PRO; Tecan) using 530/590-nm excitation/emission filters. Fluorescence reads were then normalized to the leaf weight. Superoxide anion radical accumulation was detected by NitroBlue Tetrazolium chloride (NBT; Sigma) as described previously (Basu et al. 2020) with some modifications. The 5^th^ or 7^th^ leaf was weighed and stained with freshly made NBT solution (1.0 mg mL NBT in 10 mM NaN_3_ and 10 mM phosphate buffer, pH 7.5) for 1 h. NBT-stained samples were then ground and used to quantify the generation of formazan by spectrophotometric analysis at 700 nm using a 96-well microplate reader (Infinite 200 PRO; Tecan) in case of data for *MSL10g* lines with lesions, whereas as *MSL10* overexpression lines were all analyzed using BioTek PowerWave XS2 microplate spectrophotometer (BioTek). Absorbance reads were then normalized to the leaf weight.

### Quantification of cell death

Cell death was visualized in leaves of three to five-week-old soil-grown Arabidopsis plants using Trypan blue staining. Whole plant images of *MSL10* overexpression lines were obtained with an Olympus SZx7 Stereomicroscope with DP71 digital camera and fitted with DF PL −.5X-4 objective, while Trypan blue-stained lesions were imaged using a BX53 Olympus microscope with DP80 camera digital camera fitted with a UPlanFL N 10x/0.30 objective. The size of Trypan blue-stained regions was quantified using ImageJ software as described (Fernández-Bautista et al. 2016; Basu et al. 2020). Cell death in tobacco leaves was determined by staining with FDA (500 μg/mL) and PI (1.25 μg/mL) for 15 min as described in (Veley et al. 2014). The percentage of dead cells was calculated from randomly selected fields from at least three leaves per infiltration.

### Genotyping

DNA was extracted by grinding tissue in extraction buffer (200 mM Tris-HCl pH 7.5, 250 mM NaCl, 250 mM EDTA, 0.5% SDS) and precipitating with an equal volume of isopropanol. PCR-based genotyping of *msl10-3G, msl10-1* and *msl9-1* alleles was performed as described in (Haswell et al. 2008; Basu et al. 2020).

### Immunodetection

Total proteins were extracted from leaf samples as described previously (Basu et al. 2020). Non-chlorotic leaves of 3-5-week-old transgenic Arabidopsis plants expressing MSL10-GFP or *N. benthamiana* leaves one week after infiltration with MSL10-GFP were collected. Tissue was homogenized in 2X sample buffer followed by centrifugation at 10,000g for 5 min. The supernatant was resolved by 10% SDS-PAGE and transferred to PVDF or polyvinylidene difluoride membranes (Millipore) for 16 h at 100 mA. Transferred proteins were probed with anti-GFP (Takara Bio, 1:5000 dilution; Catalog # 632380) or anti-tubulin (Sigma, 1:20,000 dilution; Catalog # T5168) as primary antibodies with incubation for overnight and 2 h, respectively. In some cases, equal loading was demonstrated by staining the PVDF membrane with Coomassie Brilliant Blue R 250 (Sigma) post immunodetection. Immunodetection was performed by incubating the immunoblot for 2 h with anti-mouse IgG conjugated with horseradish peroxidase (1:10,000 dilution; Millipore; Catalog # 12-349). Detection was performed using the SuperSignal West Dura Detection Kit (Thermo Fisher Scientific).

### Bimolecular Fluorescence Complementation (BiFC) assays

Entry vectors containing various truncated versions of *MSL10* coding region were recombined into the binary vector pDEST-VYCE(R)GW or pDEST-VYNE(R)GW (Gehl et al. 2009), which carry the C-terminal or N-terminal fragment of Venus YFP, respectively, using LR Clonase. The PCR primers used are listed in **Supplemental Table S1**. MSL10 was tagged at the C-terminus, whereas SGT1b, RAR1 and HSP90.1 were tagged at the N-terminus. For subcloning, the open reading frames (ORFs) of *SGT1b, RAR1* and *HSP90.1* were amplified from leaf cDNA using CloneAmp HiFi PCR Premix (Clonetech) and cloned into pENTR/D-TOPO (Life Technologies). Next, these ORFs were inserted into the BiFC destination vector pDEST-VYNE(R)GW using the Gateway LR Clonase II enzyme mix (Life Technologies). For MSL10, we used constructs previously generated in (Basu et al., 2020). The plasmids were introduced into *Agrobacterium strain GV3101* and pairwise combinations were co-infiltrated into 4-to-6-week-old *N. benthamiana* leaves as described (Waadt and Kudla 2008). To suppress posttranscriptional gene silencing, each construct pair was co-infiltrated with *Agrobacterium* strain *AGL-1* harboring *p19* (Voinnet et al. 1999). Infiltrated abaxial leaf areas were examined for YFP signal using a confocal microscope (Olympus Fluoview FV 3000) at 3 to 5 d post-inoculation using excitation at 488 nm and collecting emission from 495-550. The experiments were performed at least three times using different batches of plants; for each independent experiment, three *N. benthamiana* plants were infiltrated.

### Split-Ubiquitin Assay (mbSUS)

Determination of protein-protein interactions by the mbSUS assay was conducted as described previously (Obrdlik et al. 2004; Lee et al. 2019; Basu et al. 2020).The coding regions of *SGT1b, RAR1* and *HSP90.1* cloned into the BiFC destination vectors described above were used as template for amplification using universal primers as described in (Obrdlik et al., 2004). For MSL10, the construct used was described in (Basu et al., 2020). The primers used are listed in **Supplemental Table S1.** These PCR products were cotransformed with either *pMetYCgate* or *pXNGate21-3HA* yeast vectors into THY.AP4 or THY.AP5 yeast strains, respectively. All yeast vectors and strains were obtained from the Arabidopsis Biological Resource Center. Strength of interaction was quantified by β-galactosidase activity assays as described (Zhou et al. 2015).

### *Pseudomonas* infection assays

Bacterial strains *P. syringae* DC3000, *P. syringae COR^-^* (DB29; *cmaA cfa6;* (Brooks et al. 2004)) and the *hrpL* mutant (Zwiesler Vollick et al. 2002) were donated by B.N. Kunkel (Department of Biology, Washington University, St Louis, MO)*. P. syringae* bacterial strains were cultured on King’s B media plates supplemented with 50 mg mL^-1^ rifampicin for *Pto* DC3000 and 50 mg mL^-1^ kanamycin for *Pst COR^-^*. For *P. syringae* DC 3000 and *hrpL* mutant growth assays, syringe infiltration was performed as described in (Liu et al. 2015). Briefly, 10^5^ colony-forming units (CFU)/mL (OD_600_ = 0.0002) in 10 mM MgCl_2_ were injected into the underside of the leaves of 5-week-old plants using a needleless 1 cc syringe. For *P. syringae COR^-^*, spray-inoculation was performed at a bacterial titer of OD_600_ of 0.2, which corresponds to the final inoculum concentration of 1 × 10 CFU (Jacob et al. 2017). After the indicated days of infection, leaf discs were obtained from three separate plants belonging to each genotype per bacterial strain tested. Bacterial titer was determined by grinding leaf discs to homogeneity in 10 mM MgCl_2_ using a bead mill and plating a dilution series onto KB plates with appropriate selection. Colonies were counted and used to calculate the mean CFU/cm for each treatment, and the values were log-transformed. The log-transformed means from individual biological replicates, as single data points, were then combined from three to five independent biological trials and used to calculate the mean and standard error.

### Virus-Induced Gene Silencing (VIGS) and Agrobacterium-mediated transient assays

Plasmids pTRV1 and pTRV2 were used for VIGS assay as previously described (Voinnet et al. 1999). Constructs pTRV2:00, *pTRV2:NbSGT1*, pTRV2*:NbRAR1*, and *pTRV2:LeHSP90* were obtained from the Arabidopsis Biological Resource Center. The empty vector, *pTRV2:00* was used as a negative control. pTRV2*-NbPDS (PHYTOENE DESATURASE)* was used as a positive control for silencing. All these constructs were transformed into *A. tumefaciens* strain *GV3101*. For leaf infiltration *GV3101* harboring pTRV1 or transformed with one of the pTRV2 constructs were mixed at a 1:1 ratio to a final OD_600_ of 0.5 and then co-infiltrated into 2-week-old *N. benthamiana* leaves. Silencing efficiency was evaluated nine days after infiltration by performing RT-qPCR analysis. After confirming suppression of *SGT1, RAR1* and *HSP90*, infiltration of *MSL10-GFP, MSL10^4A^-GFP* and *MSL10^4D^-GFP* was performed to a final OD_600_ of 0.5 along with *p19* to a final OD_600_ of 0.3, as in (Maksaev et al. 2018). Each silencing experiment was repeated three times, and each experiment included at least three different plants.

### Stomatal aperture

Plants were grown under short-day conditions for 5-6 weeks. Intact leaves (5^th^, 7^th^ or 9^th^) were used for aperture measurements as described (Eisele et al. 2016). Leaves were exposed to white light for 1 hour while floating (abaxial side touching the solution) in stomatal incubation buffer containing 10 mM MES-KOH, 10 mM KCl, 50 μM CaCl_2_, pH 6.5. Subsequently, one-half of leaf was immersed in stomatal incubation buffer with the bacterial suspension at 1×10^8^ CFU/ml, OD_600_Ū=Ū0.2 and incubated for 4 h; whereas the other half of the leaf was incubated in water. Abaxial leaf surfaces were imaged, and stomatal apertures measured with ImageJ as described (Chitrakar and Melotto 2010). To monitor stomatal closing in response to ABA, leaf discs with pre-opened stomates were exposed to the same buffer supplemented with 10 μM ABA for a 2 h under the same conditions used in (An et al. 2016). Independent trials of the experiments were done around the same time of the day for each of the independent trials and the treatments to avoid potential rhythmic effects of photoperiod on stomatal aperture.

### Confocal Microscopy

The subcellular localization of MSL10-GFP fusion proteins in epidermal cells of *N. benthamiana* leaves and Arabidopsis, as well as FDA/PI dual stained tobacco epidermal cells, were visualized using on an inverted FV3000 (Olympus) confocal microscope using a 20X objective. The percentage of dead cells was quantified as outlined in (Veley et al. 2014; Maksaev et al. 2018). FDA and PI dual-stained epidermal cells were imaged sequentially using the 488-nm laser to excite FDA and the 561-nm laser to excite PI. Emission was detected between 505-525 nm and between 580-660 nm for FDA and PI, respectively. The setting used for capturing FDA signals was also used for detecting GFP signal.

### Statistical analyses

Statistical evaluations for all figures were conducted using R Studio software (RStudio) and GraphPad Prism version 7.0. Statistical differences were analyzed as indicated in the figure legends. Tukey’s HSD post-hoc test was used to determine statistical significance for balanced data sets. Scheffe’s post-hoc test was used to determine statistical significance for unbalanced data sets.

### Accession Numbers

Sequence data from this article can be found in the Arabidopsis Genome Initiative database or GenBank under the following accession numbers: *MSL9* (At5G19520), *MSL10* (At5G12080), *SAG12* (At3g20770), *PERX34* (At3g49120), *DOX1* (At3g01420), *OSM34* (At4G11650), *UBQ5* (At3g62250), *EF1α* (At5g60390), *PAD4* (At3g52430), *SID2* (At1g74710), *SGT1b* (At4g11260), *HSP90.1* (At5g52640), *RAR1* (At5g51700), *EDS1* (At3g48090), *NDR1* (At3g20600), *PR1* (At2g14610).

## AUTHOR CONTRIBUTIONS

B. and E.S.H. conceived the project and designed the experiments. J.M.C. contributed Figure 4.; K.M.V contributed data to Figure 1, D.B. performed the rest of the experiments. D.B. and S.H. wrote the manuscript with input from J.M.C and K.M.V.

## ACKNOWLEDGEMENTS

We thank Barbara Kunkel (Washington University in St Louis) for providing *pad4* mutant seeds and all *Pseudomonas* pathovars. All other mutant seeds were obtained from the Arabidopsis Biological Resource Center. This work was supported by NSF MCB-1253103 and the NSF Center for Engineering Mechanobiology Award 1548571. J.M.C. was supported by NSF Graduate Research Fellowship DGE-1745038.

## SUPPLEMENTARY FIGURES

Figure S1. Growth at high temperatures suppresses dwarfing associated with the *msl10-3G* allele and expression of the *MSL10g^7A^* transgene.

Figure S2. Transcript and MSL10-GFP abundance in response to VIGS.

Figure S3. Validation of the *sgt1b-2* allele and phenotypic characterization of *sgt1b-1, sgt1b-2, rar1-21* and *hsp90.1* mutants expressing MSL10-GFP.

Figure S4. MSL10 does not interact with SGT1b, RAR1, or HSP90.1 in yeast two-hybrid or BiFC assays.

Figure S5. Cell death phenotypes after transfer from high to low temperatures.

Table S1. Oligonucleotides used in this study.

