## Supplemental Data for "The Mechanosensitive Ion Channel MSL10 Modulates Susceptibility to *Pseudomonas syringae* in *Arabidopsis thaliana*"

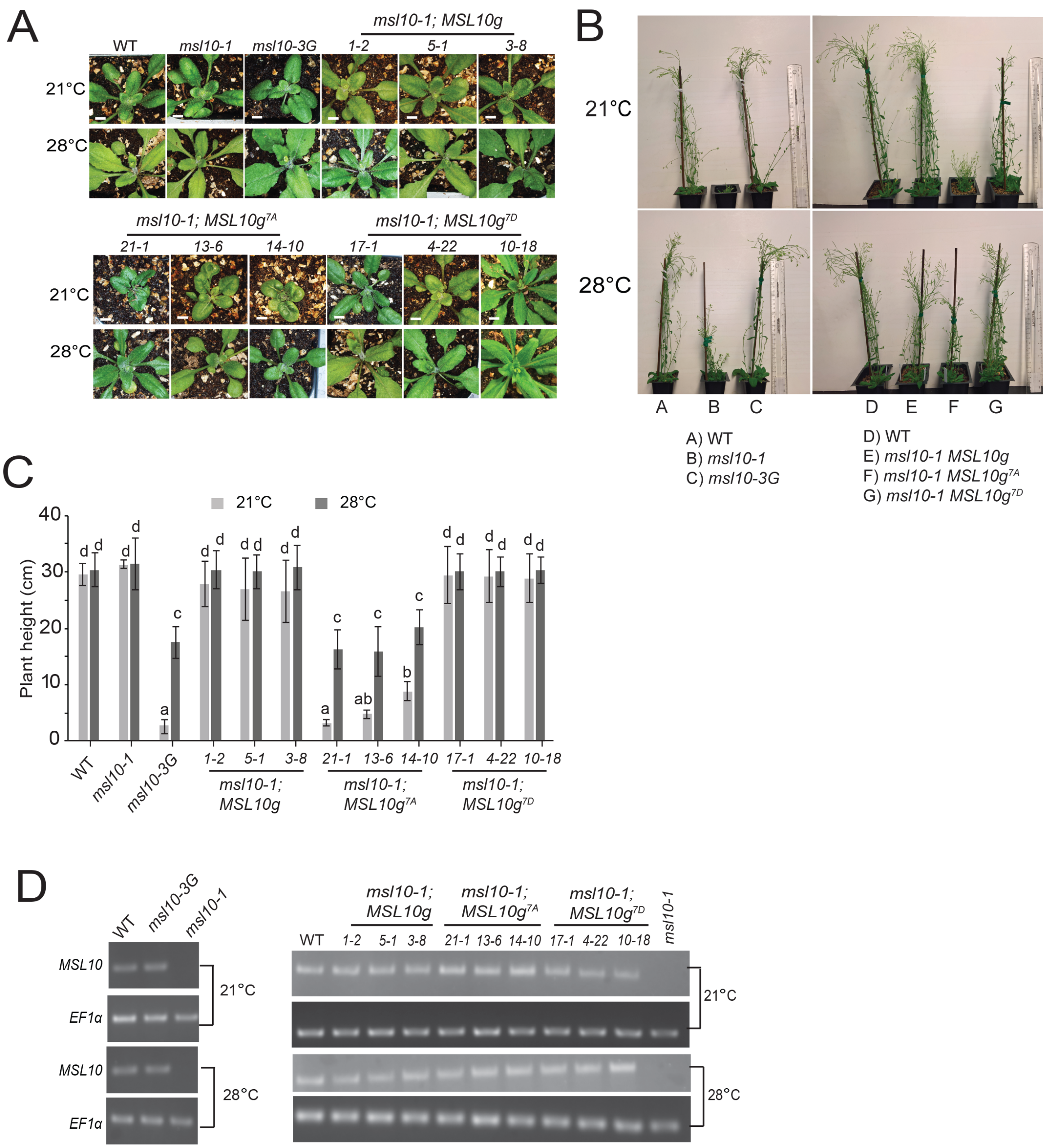


**Figure S1. Growth at high temperatures suppresses dwarfing associated with the *msl10-3G* allele and expression of the *MSL10g*$^{7A}$ transgene.** (A) Images of four-week-old plants from wild type, *msl10-1* and *msl10-3G* mutants, as well as individuals from three independent T4 homozygous lines expressing MSL10-GFP phospho-variants, grown side-by-side on soil under 24 h light at either 21°C or 28°C. Bar, 0.5 cm. (B) Eight-week-old plants of the genotypes listed in (A). (C) Plant height of the indicated genotypes. Data are means ± SD of three independent trials, each with 15-18 plants. Different letters indicate significant differences, as determined by two-way ANOVA followed by Tukey's post-hoc test ($P < 0.05$). (D) Semi-quantitative RT-PCR amplification of *MSL10* transcripts from the indicated genotypes. cDNA was synthesized from total RNA extracted from three-week-old plants grown as in (A). *EF1α* (Elongation factor 1 alpha) was used as a loading control.

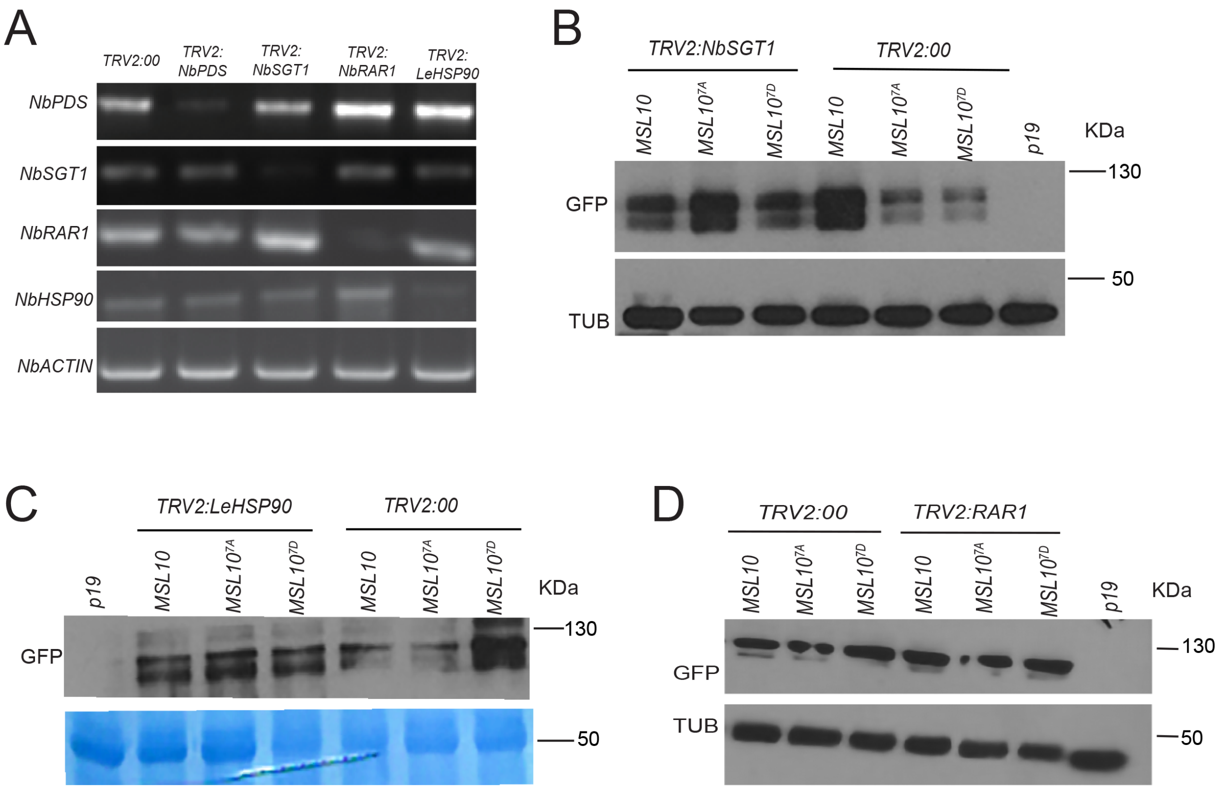


**Figure S2. Transcript and MSL10-GFP abundance in response to VIGS.** (A) *N. benthamiana* leaves were infiltrated with Agrobacterium carrying *TRV1* and *TRV2-NbSGT1, TRV2-NbRAR1, TRV2-LeHSP90, TRV2-PDS* or *TRV2* alone *(TRV2:00*). RNA was extracted from leaves 10 days after infiltration and *PDS, SGT1b, RAR1,* and *HSP90* gene expression analyzed by qRT-PCR. *NbACTIN* transcript levels are shown as a loading control. (B-D) Immunoblot of MSL10-GFP protein levels in (B) TRV2-LeHSP90- (C) TRV2-NbSGT1- and (D) TRV2-NbRAR1-infected *N. benthamiana* leaves 10 days after infiltration. Total protein extracted from leaves was analyzed by SDS-PAGE followed by immunoblot analysis using anti-GFP antibodies. Coomassie Brilliant Blue staining of the blot or immunoblot analysis with anti-alpha-tubulin served as loading control.

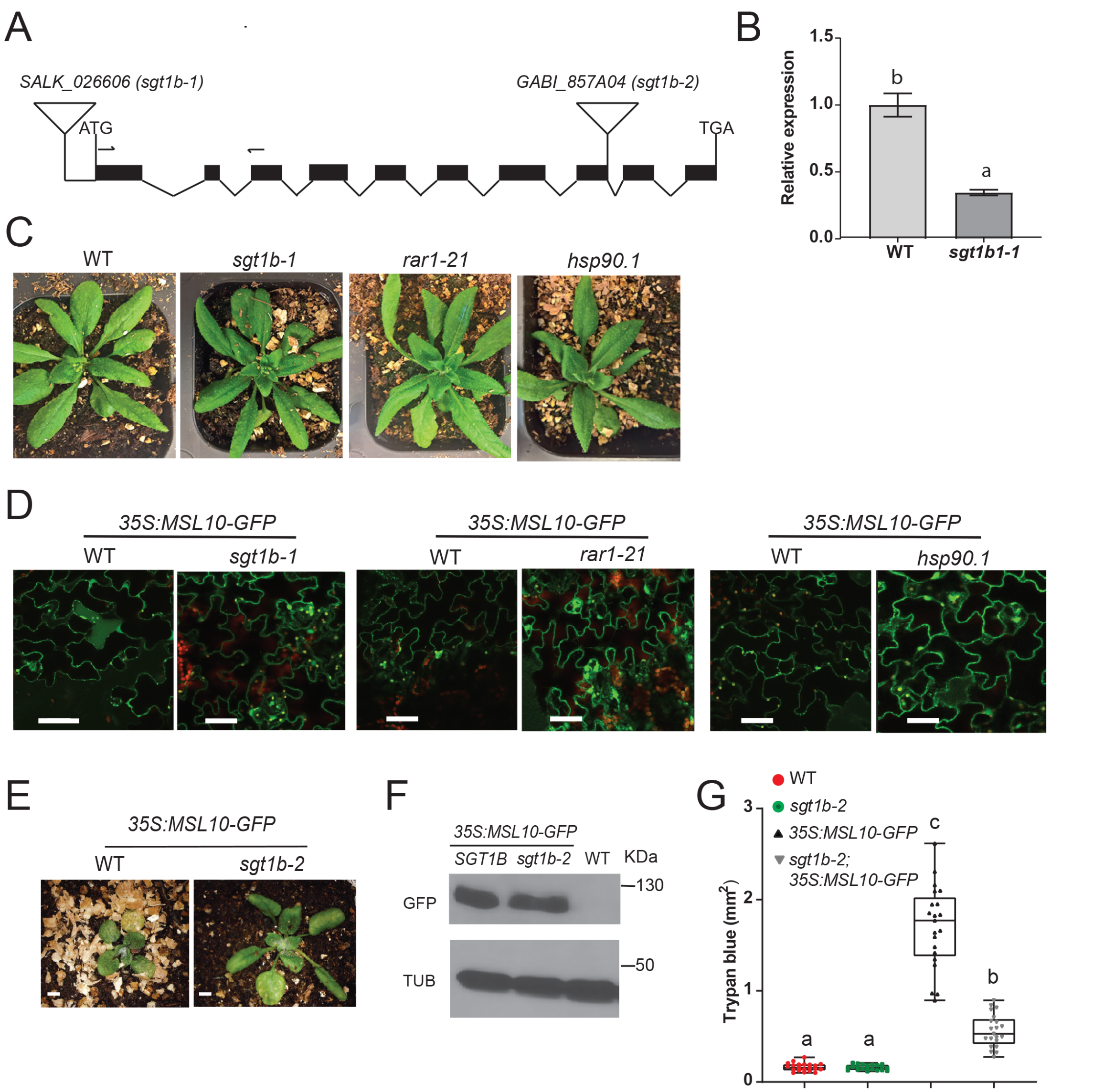


**Figure S3. Validation of the *sgt1b-2* allele and phenotypic characterization of *sgt1b-1, sgt1b-2, rar1-21* and *hsp90.1* mutants expressing MSL10-GFP.** (A) Diagram of At4g11260 (*SGT1b*) and the T-DNA insertion sites in SALK_026606 and GABI_857A04 lines. The black boxes indicate exons and the lines in between represent introns. Inverted triangles and arrows indicate T-DNA insertions and the positions of genotyping primers, respectively. (B) Relative transcript levels of *SGT1b* in rosette leaves of wild-type and *sgt1b-2* mutant lines. RNA was extracted from four-week-old plants grown at 21°C under a 24 h light regime, and expression of *SGT1b* transcripts measured by qPCR as described in Figure S1. (C) Representative images of five-week-old *sgt1b-1, rar1-21, hsp90.1* and wild-type plants grown side-by-side on soil under 24 h light at 21°C. (D) Representative confocal images of GFP signal at the periphery of epidermal cells from four-week-old *35S:MSL10-GFP* siblings segregating *sgt1b-1, rar1-21*, or *hsp90.1*. Plants were grown at 21°C under a 24 h light regime. Chlorophyll autofluorescence is shown in red. Bar, 20 μm. (E) Representative images of plants expressing MSL10-GFP in *SGT1b* or *sgt1b-2* backgrounds. Three-week-old plants were grown at 21°C under a 24 h light regime. Bar, 0.5 cm. (F) Immunoblot showing MSL10-GFP levels in wild-type plants and siblings from *35S:MSL10-GFP* lines segregating *sgt1b-2*. Total protein was extracted from three-week-old rosette leaves grown at 21°C under 24 h of light. (G) Trypan blue staining in wild type and *sgt1b-2* mutant rosette leaves. Three-week-old plants of indicated genotypes were grown under 24 h light at 21°C. Data represent the means ± SD from three independent trials, each trial with 7 leaves. Different letters indicate significant differences as determined by one-way ANOVA followed by Tukey's post-hoc test ($P < 0.05$).

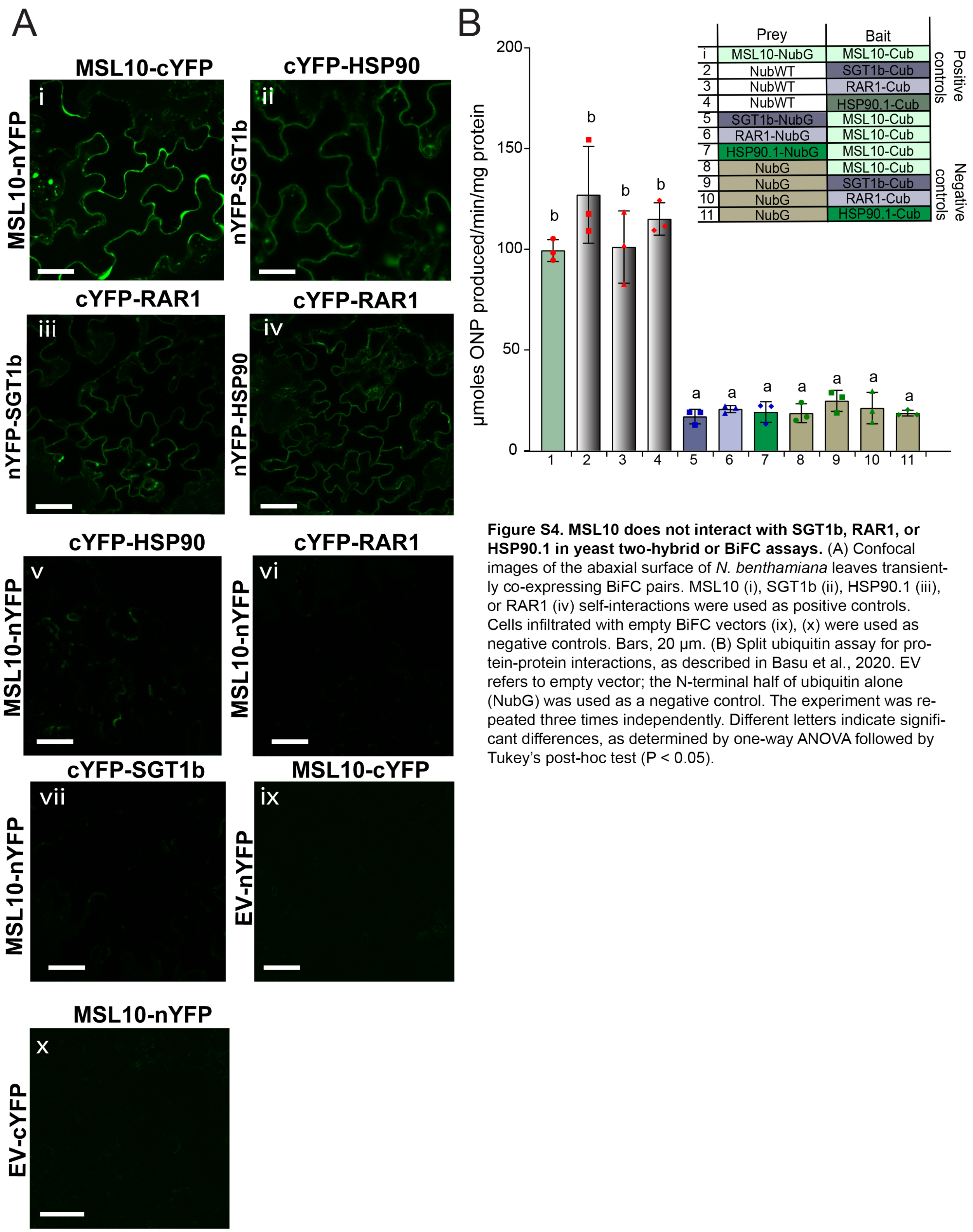


**Figure S4. MSL10 does not interact with SGT1b, RAR1, or HSP90.1 in yeast two-hybrid or BiFC assays.** (A) Confocal images of the abaxial surface of *N. benthamiana* leaves transiently co-expressing BiFC pairs. MSL10 (i), SGT1b (ii), HSP90.1 (iii), or RAR1 (iv) self-interactions were used as positive controls. Cells infiltrated with empty BiFC vectors (ix), (x) were used as negative controls. Bars, 20 µm. (B) Split ubiquitin assay for protein-protein interactions, as described in Basu et al., 2020. EV refers to empty vector; the N-terminal half of ubiquitin alone (NubG) was used as a negative control. The experiment was repeated three times independently. Different letters indicate significant differences, as determined by one-way ANOVA followed by Tukey's post-hoc test ($P < 0.05$).

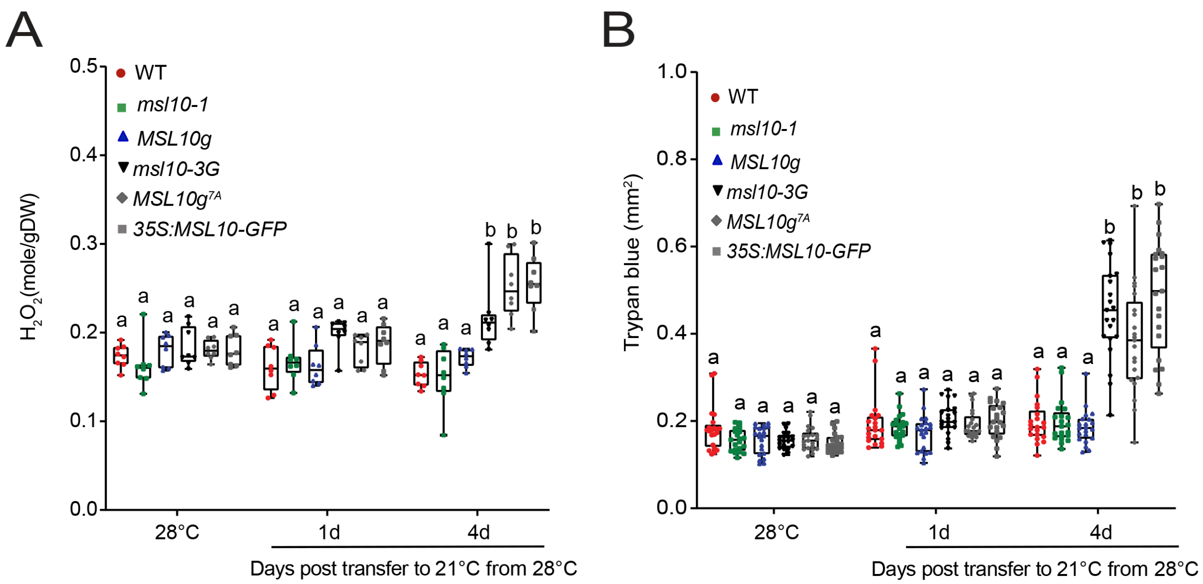


**Figure S5. Cell death phenotypes after transfer from high to regular temperatures.** (A) Plants of the indicated genotypes were grown for five to six weeks at 28°C under 10 h of light/14 h of dark. Three-week-old rosette leaves were collected for quantification of hydrogen peroxide ($H_2O_2$) from plants transferred to 21°C for 1 or 4 days. using the Amplex Red-coupled fluorescence quantitative assay. Data represent the means ± SD from three independent trials, each with 3-4 leaves. $H_2O_2$ content was quantified as μmol $H_2O_2$ per mg dry weight (DW). (B) Quantification of cell death detected by Trypan blue staining in rosette leaves as in (A). Data represent the means ± SD from three independent trials, each with 7 leaves. Different letters indicate significant differences as determined by two-way ANOVA followed by Tukey's post-hoc test ($P < 0.05$).

**Supplemental Table 1.** Oligonucleotides used in this study.

| Name | Primer 5'-3' |
|---|---|
| **BiFC constructs** | |
| *SGT1bF* | CACCATGGCCAAGGAATTAGCAGAGA |
| *SGT1bR* | CCACTTCTTGAGCTCCA |
| *RAR1F* | CACCATGGAAGTAGGATCTGCAACGAAGAAGC |
| *RAR1R* | GACCGCCGGATCAGGGCTGCTGCTA |
| *HSP90.1F* | CACCATGGCGGATGTTCAGATGGCTGATGC |
| *HSP90.1R* | GTCGACTTCCTCCATCTTGCTCTCTTC |
| **Split ubiquitin yeast two-hybrid constructs** | |
| *Universal F (attBF1)* | ACAAGTTTGTACAAAAAAGCAGGCTCTCCAACC ACCATG |
| *Universal R (attBR1)* | TCCGCCACCACCAACCACTTTGTACAAGAAAGC TGGGTA |
| **RT-PCR and qRT-PCR** | |
| *SGT1bF* | ATGGCCAAGGAATTAGCAGAG |
| *SGT1bR* | TTCTTCTGCAATACGAAGATCG |
| *PR1F* | ATCGGTTTCTTCTTCTTATCGTG |
| *PR1R* | TTTGATTTCTGATCGACGGGG |
| *MSL10 F* | AGAGGTTGATCTTGTGTTCCC |
| *MSL10 R* | TTTGTGTCGTTTAAGGAATGCG |
| *UBQ5 F* | TCTCCGTGGTGGTGCTAAG |
| *UBQ5 R* | GAACCTTTCCAGATCCATCG |
| *EF1alpha F* | ACAGGCGTTCTGGTAAGGAG |
| *EF1alpha R* | CCTTCTTCACTGCAGCCTTG |
| *MSL10 F* | GTTGGTTTCTGGGTTTAAGCC |
| *MSL10 R* | TACTTGGAGTAACCGGTGCTG |
| *NbPDSF* | TCCTCACGCCCAACTAAACCATT |
| *NbPDSR* | CTTCAACATAAGATTGCCCTCCA |
| *NbSGT1F* | AGGCCAACATCAAACTCAAC |
| *NbSGT1R* | CAAACAACCTAGGCTGGAAA |
| *NbRAR1F* | CCACCTTCACTGAGGATGAT |
| *NbRAR1R* | AAAGGTCTTACCACAGCCCT |
| *NbHSP90F* | GATCCTGAAGGTTATTCGCA |
| *NbHSP90F* | CAAGCTTGAGACCTTCCTTG |
| *NbEDS1F* | TGTTGGCACAGATGAGGTAGCCA |
| *NbEDS1R* | CCCGACGAGTGCCCTGCAAA |
| *NbACTIN4F* | TGGGTTTGCTGGAGATGATG |
